# Sparrowhawk: harnessing WebAssembly for private microbial genomics, and beyond

**DOI:** 10.64898/2026.11.18.767565

**Authors:** Víctor Rodríguez-Bouza, Johanna H. von Wachsmann, Bede Constantinides, Antoine Andréoletti, Erkison E. Odih, Luca Nesterenko, Vikas Gupta, John A. Lees

**Affiliations:** EMBL’s European Bioinformatics Institute (EMBL-EBI); University of Birmingham; Université de Montpellier; London School of Hygiene and Tropical Medicine (LSHTM); Université Claude Bernard Lyon 1 (UCBL)

**Keywords:** bacterial genomics, genome assembly, phylogenetics, antimicrobial resistance, WebAssembly, privacy

## Abstract

Computational methods are a growing component of every scientific discipline, but the pace of training and expertise for using these methods has not kept up with their widening usage. Researchers expect state-of-the-art methods to be able to independently explore their results, and do so dynamically and interactively. Producing such an application which implements a new algorithm, even though it would access a much wider audience, is typically beyond the expertise and time constraints of computational method developers. Here, we show our `sparrowhawk` framework can rapidly create an interactive web application to run interactive analysis on a user’s machine, across a wide range of algorithms.

Focusing first on genomic epidemiology, a domain with a specific need for such methods, we created novel lightweight methods for genome assembly, antimicrobial resistance gene annotation, and taxonomic identification. Our results exceed performance of previous lightweight methods, and produce results that reproduce or closely match previous analysis pipelines. By using WebAssembly, our framework benefits from not requiring installation, command line knowledge, or transferring sensitive data over an internet connection.

We show how algorithms from other researchers, and spanning fields from machine learning to physics, can rapidly be ported to web tools using our framework. We demonstrate an immediate impact on microbial bioinformatics, but expect that uptake of `sparrowhawk` by developers of other tools broadly will help other researchers, and create new tools for teaching and public engagement. Thanks to the broad availability and familiarity with web browser tools across society, `sparrowhawk` has the potential to significantly lower the barrier to entry to advanced computational methods.

---

Data science and statistics are key elements of modern biomedical research, but there is a gap between researchers with domain expertise collecting data, and specialists developing complex methods to analyse this data. Many generators and consumers of this data would benefit from being able to analyse it themselves – they have a unique knowledge of biases and important questions from their data, and if they are able to analyse the data locally they can answer these questions more quickly.

Existing platforms allow people not experienced in command line interface usage to perform some bioin-formatics analyses steps more easily. Examples include phylogenetic studies, multiple sequence alignment, or antimicrobial resistance (AMR) genes detection, offered by platforms such as Job Dispatcher [1], the online bioinformatics tools from the Center for Genomic Epidemiology of DTU [2, 3], or the Galaxy online software [4]. These platforms work in a similar fashion: they require the user to provide the input data for the method to be used through a simple and intuitive interface. This data is uploaded to remote servers and is usually processed in a high performance computing (HPC) cluster, and after processing is complete the user receives a notification that the outputs can be retrieved. However this approach has drawbacks: it requires the user to upload the data to a remote server, which can pose an issue if the data is sensitive [5] or there are data sovereignty concerns [6]; the process itself is reliant on the maintenance of remote computational servers; analysis cannot easily be altered and explored interactively with intuitive visuals.

Inspiration for how to empower biomedical researchers with computational methods can be taken from some of the most successful data analysis packages, Excel and RStudio. Both offer easy installation of a single package which works on many different computers. Visualisation of data and the results of methods is directly integrated into the tools. All data is analysed on the user’s computer, which negates issues of sharing private data or needing an internet connection to upload data for analysis elsewhere. Interaction with the data and methods is dynamic, letting users try different options and see how they affect the results – and in doing so can often gain a deeper understanding of their own data and how different methods work. When stuck, users can find ample documentation and examples.

A potential bridge between complex methods and interactive visualisation is WebAssembly. By running in a web browser on a user’s computer, WebAssembly gives the advantages of software like Excel, without compromising data privacy. Complex methods can run quickly in a web browser, without requiring method developers to rewrite their actual algorithms or code. Examples of WebAssembly usage in the biological fields exist already [7–10], and also some examples in other fields, such as astrophysics [11] or scientific visualisation [12]. However, research uses of WebAssembly are relatively few in number, documentation is limited, and requires coordination of data across multiple programming languages.

In this manuscript we present `sparrowhawk` (figure 1): a suite of novel and adapted methodologies presented as web platform. `sparrowhawk` uses WebAssembly [13–16]: a binary file standard designed for creating executables that run on Internet browsers. We first report the design of novel, lightweight algorithms for genome assembly, antimicrobial resistance gene identification and taxonomic identification that are adapted to the constraints of this platform, showing results which match those from standard tools but with greatly reduced resource use. We then apply our technique directly to a range of fundamental methods in public health genomics and metagenomics, which both due to training gaps and the need for rapid turnaround and exploration of results have a clear need for accessible methods [17]. We create further tools for mapping reads, phylogenetic tree reconstruction, host read removal, and protein sequence embedding using a transformer-based machine learning model.

**Fig 1.**
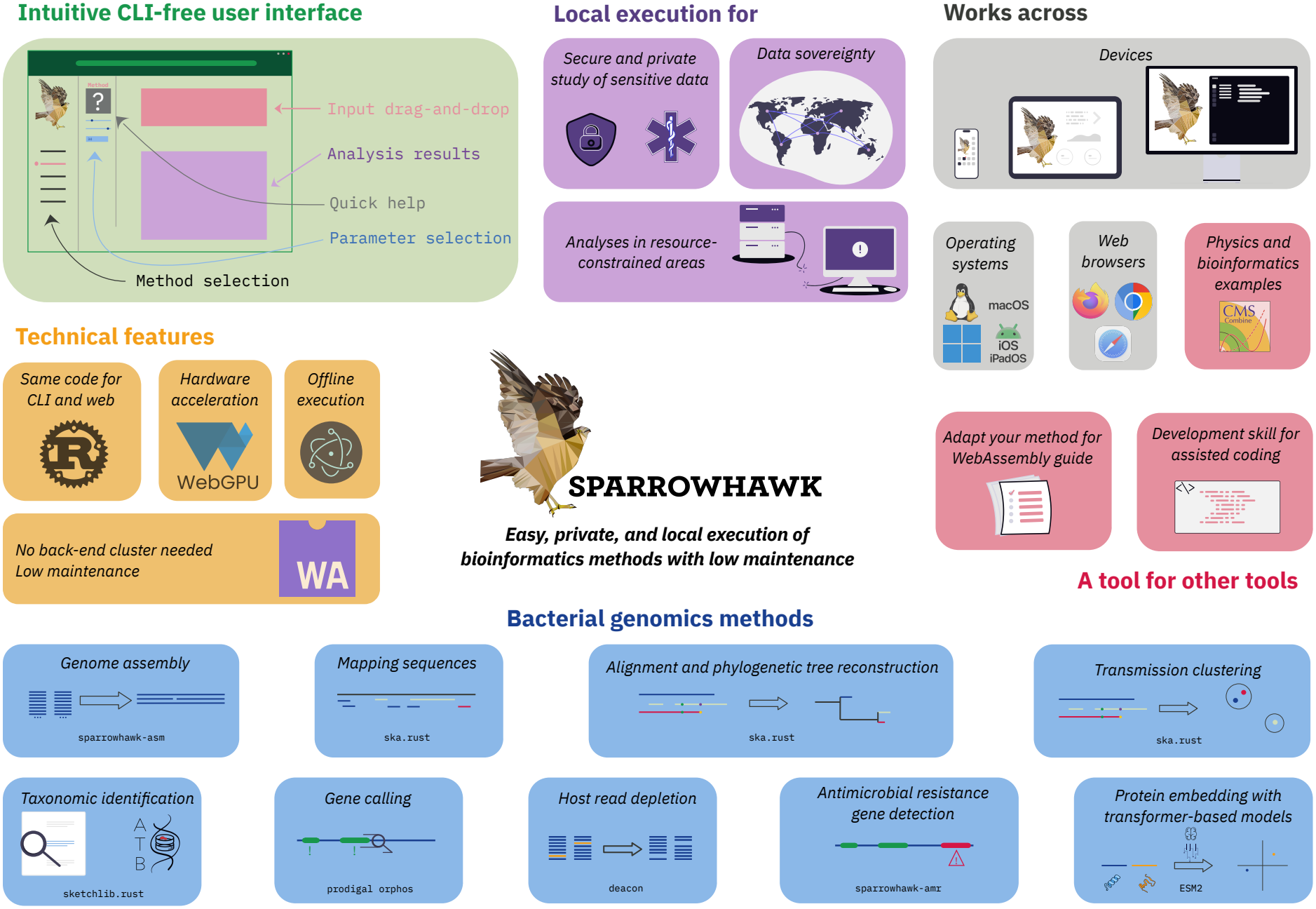
Overview of the `sparrowhawk` platform for bacterial genomic analyses. `sparrowhawk` provides a suite of bacterial genomics analyses through a simple and intuitive web interface that can run in any major Internet browser, operating system, across devices (computer, tablet, smartphones). Methods can share execution code so do not need to be separately maintained. By using the open standard WebGPU, we can provide additional hardware acceleration that especially benefits complex machine learning models. Any analysis done with `sparrowhawk` only needs to access the website to be ready for running: all computation is local, so no input data upload is needed. This helps particularly with sensitive data, resource-constrained environments, or with data sovereignty or legal concerns. `sparrowhawk` lowers the barrier for other method developers to adapt their algorithms for local execution on a web browser, by offering a guide on how to design an interface and avoid the most common errors. We also provide a development skill designed for automated agentic development, that we have refined using our experience on adapting bioinformatics and particle physics methodologies.

Running on the user’s device removes the need for input data upload and output download, and thus the need to transfer large files over the network, helping in those situations when the bottleneck can be the Internet connection. Also, the data does not leave the user’s system, thus preserving privacy and data sovereignty. By using compiled code the methods achieve good computational performance, and relatively easy acceleration using the client’s GPU through WebGPU. Finally, the portability by design that WebAssembly has allows `sparrowhawk` to work with different Internet browsers and different devices (e.g. smartphones, tablets), adapting to a myriad of different needs, situations, and environments.

In addition to a genomic epidemiology package, `sparrowhawk` can also be used by other tool developers to create secure web applications implementing their methods, without any need to maintain a back-end server. We have created a guide to develop new methods or adapt existing ones designed to work with WebAssembly, showing how to solve common issues. We also create skills which support autonomous adaptation of existing methods using AI agents, which we demonstrate on an external read aligner (`rammap`, based on `minimap2` [18, 19]), an embedding and clustering tool (`Mandrake` [20]), and a high-energy physics application (the particle collision data statistical analysis tool CMS Combine [21]). `sparrowhawk` is accessible at https://sparrowhawk.bacpop.org.

## Results

### Novel lightweight algorithms for bacterial genomics

Bioinformatic analysis is usually targeted at a high-performance computing environment, with comparatively abundant computational resources. Most algorithm development therefore tends to focus on maximising performance and/or minimising CPU time so that larger analyses can be run in a fixed budget. Running within a web environment imposes constraints on disk, memory and CPU use that must be a priority consideration to develop a responsive interactive tool. While some approaches natively fit these constraints, some of the more computationally demanding fundamental methods do not. Typically this is because of high memory requirements, or the need for large external databases. We therefore first describe our development and benchmarking of lightweight algorithms for sequence assembly and taxonomic identification.

As genome assembly is the first step of most bacterial genomic analyses, we developed `sparrowhawk-asm`, a short-read assembler designed to compile to WebAssembly. Aimed at short reads, which make of the majority of sequencing runs uploaded to public repositories (*≈*93 % of all ENA submissions between 2019 and 2024 [22]), `sparrowhawk-asm` follows a traditional de Bruijn graph approach. We were inspired by the approach of `Minia`, which has been used to assemble the entire SRA [23]. Because bacterial genomes are relatively small (*≈*1 *−*10 Mbp [24]), we can be more flexible with peak memory usage, which has a much larger effect with larger genomes such as in most eukaryotes. For example, this allows loading into memory the whole set of reads to be assembled. `sparrowhawk-asm` uses this flexibility to be a simple, fast, and robust assembler by leveraging *k*-mers (subsequences of length *k*), and a rolling-hashing function for counting and neighbour search, as used in other methods [25–27]. The assembler can be used natively as a command line tool, but also as part of the `sparrowhawk` website.

We have evaluated `sparrowhawk-asm` using both real sequencing runs and simulations coming from different species (as described in Methods) and running natively. We compared our approach to `SPAdes` [28], `SKESA` [29], `Minia` [30], and `Velvet` [31], using four threads in all executions. The results show all assemblers recover a similar fraction of the original genome in simulations (*≈* 98 % for multi-*k* methods). `sparrowhawk-asm`’s assemblies showed few assembly errors overall, close to `SKESA`, which recovered the original assemblies with fewest deviations. This can be explained by a slightly more conservative approach when correcting the graph compared to `Minia`, allowing for an increased accuracy on the results while getting lower or similar contiguity results to that assembler (fig. 2b,c). Both `SPAdes` and `SKESA`, methods with a more elaborate pipeline which includes paired-end read information usage, achieve the best contiguity results.

**Fig 2.**
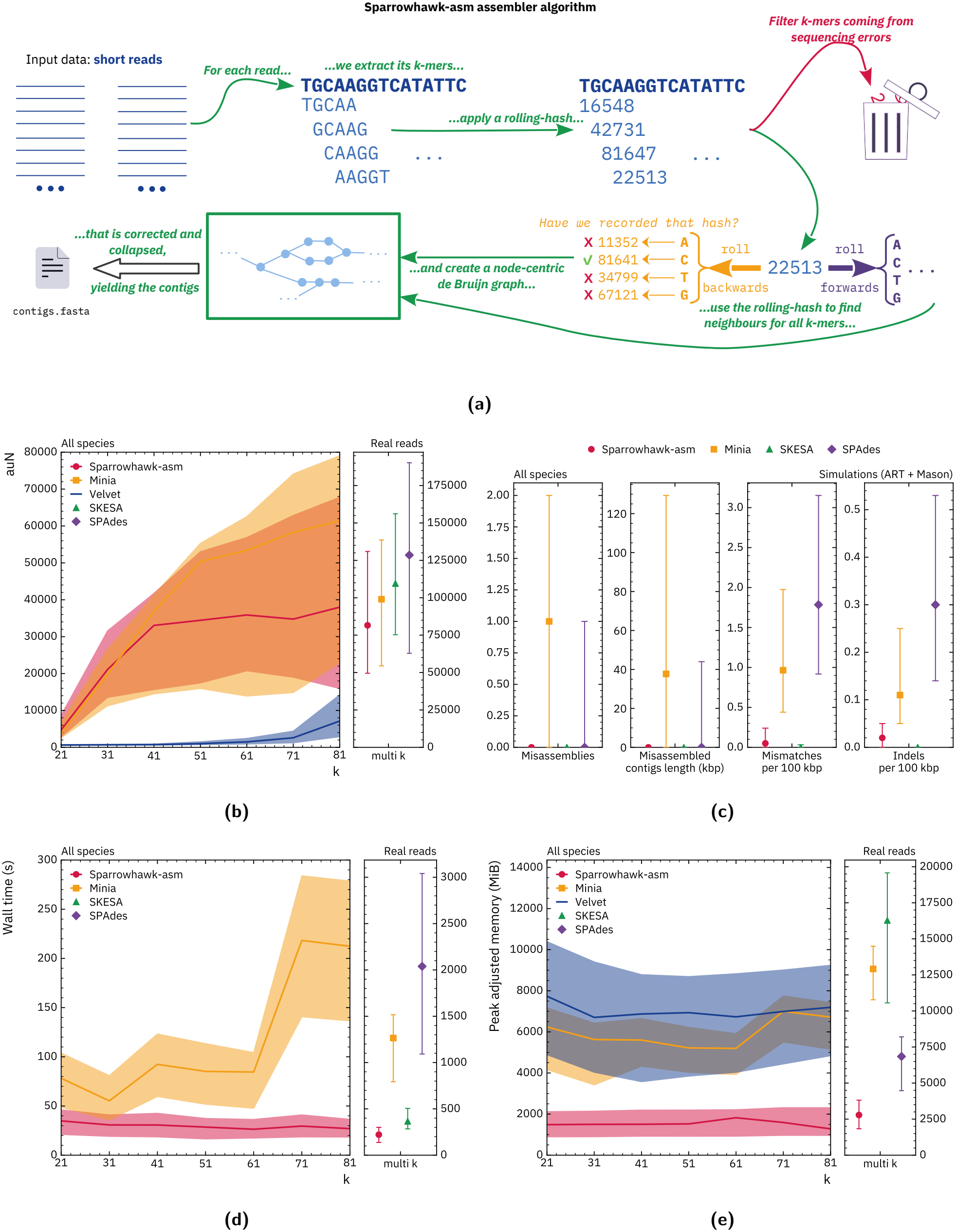
**a**, Single-*k* execution of `sparrowhawk-asm`. **b**,**c**,**d**,**e**, Evaluation of `sparrowhawk-asm` in native command-line execution on both read simulations and real data. In all plots, five assemblers are compared: `SPAdes`, `SKESA`, `Minia`, `Velvet`, and `sparrowhawk-asm`, using `Minia` and `sparrowhawk-asm` as both single-*k* and multi-*k*. Uncertainty areas show the interquartile range, and the clear lines or markers show the median of the distribution arising from the constructed simulations or the dataset of reads from real sequencing experiments. **b**, Area under the curve of the *N* function, using real sequencing data. **c**, Number of misassemblies, mismatches per 100 kbp, insertions and deletions (indels) per 100 kbp, and misassembled contigs length (in kbp) using `ART` and `Mason` simulations, for the multi-*k* runs. **d**, Total execution (wall) time with real sequencing data. `Velvet` median values ranged between *≈* 500 s and *≈* 1,100 s. **e**, Maximum memory (RSS) and filesystem allocations on disk (in the case of `SPAdes` and `Minia`) with real sequencing data.

Running inside a browser imposes a memory ceiling and slower execution than native code. Long running times inside the browser are more of an issue than jobs running on servers, as they cannot as easily be kept running without the user present. To cope, `sparrowhawk-asm` counts *k*-mers with Bloom filters and removes erroneous *k*-mers using a threshold fitted to the *k*-mer spectrum. In combination with central design choices such as using a rolling-hashing function for fast *k*-mer access, `sparrowhawk-asm` showed the lowest total execution times for single and multi-*k* assemblies, and lowest peak memory usage (peak *adjusted* memory, fig. 2). `SKESA` has a somewhat similar run time (median values of 222 s `sparrowhawk-asm` vs 365 s `SKESA`). `Minia` has a median wall time 5.6 times larger than `sparrowhawk-asm` for the multi-*k* scenario, and 1.8*−* 7.9 times larger for single-*k* runs (increasing in *k*). More detailed results can be seen in supp. figs. 1–18.

`sparrowhawk` also provides bacterial taxonomic identification, where a genome is assigned to a species identifier. Here the challenge is not memory but the reference database, which for standard classifiers is too large to download with a web page. `sparrowhawk` circumvents this issue by taking advantage of genomic sketches, an inverted index (figure 3), phylogenetic compression [32], and dereplication [33]. We take advantage of the AllTheBacteria dataset [34]: a collection of currently *≈*3 M high-quality, systematically assembled genomes from multiple bacterial species available in public repositories. We first apply a method for genome clustering that dereplicates these genomes into 7305 clusters [33], from which we extract the best quality assembly per each and create an inverted index with `sketchlib.rust`, reordering samples to make more closely related samples adjacent, to improve compressibility. To create the index, we chose *k* = 17 and *s* = 50, balancing accuracy (98 %) against its size (fig. 3). This yields fast genomic comparisons with a small 3.5 MB index.

**Fig 3.**
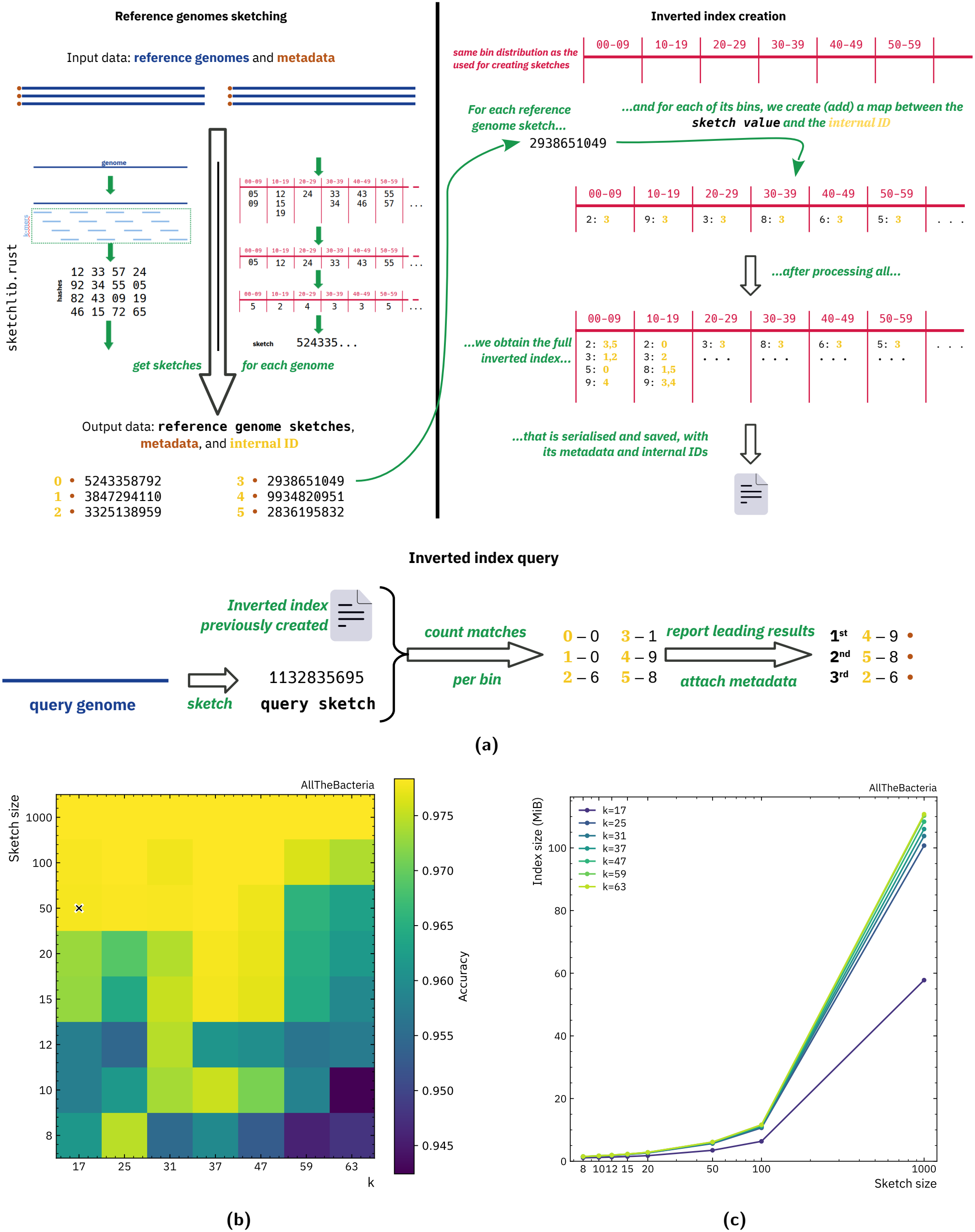
Overview of the taxonomic identification algorithm. **a**, `sparrowhawk` uses small genome sketches and an inverted index feature to create a taxonomic reference compact enough to be shipped with the website. This diagram shows the different steps to create the index. The reference genomes are sketched offline. Then, an inverted index is created from the sketches, and it is serialised and stored on disk. In the browser, query genomes are first sketched with the same parameters as the inverted index. Then, with the information stored in the index, the algorithm checks per each bit bin how many matches we have against each reference genome, here represented by their internal index identification number. Finally, we report the results of the first three ranks. **b**, Evaluation of the main parameters for inverted index construction, *k* and the sketch size *s*, using the AllTheBacteria dataset, showing as accuracy the agreement with `GTDB-Tk` labels. **c**, Index size with the sketch size for multiple values of *k*.

### Sparrowhawk can be used to rapidly identify bacterial transmission and resistance genotypes

To demonstrate that our lightweight tools provide adequate results, in a setting where rapid interactive investigation is most needed, we have reanalysed data from an international pathogen transmission study on *Klebsiella pneumoniae* [35]. In addition to the tools above, we ported to WebAssembly existing lightweight *k*-mer methods from `ska.rust` [26, 27] for variant calling, and additionally implemented neighbour-joining phylogenies and single linkage clustering based on adjustable SNP thresholds. The results – variants detected, phylogenetic tree, distances, an alignment, or SNP cluster assignments – can be downloaded from the website, or directly viewed using integrated methods such as `MSEABOARD` [36] or `Taxonium` [37].

We processed existing read data of the study using `sparrowhawk-asm` to obtain, in combination with the existing assemblies online, 1,522 assemblies that were afterwards analysed with `sparrowhawk`’s “Alignment” tab with five processing workers (22 minutes, with a peak memory usage of 6.5 GB). We then ran the transmission clustering analysis with the location and temporal metadata uploaded with default parameters (fig. 4). Supp. figs. 21–23 show how transmission analyses are displayed on the website. The number of clusters recovered is similar (138 `sparrowhawk`, 156 reference), and so is their arrangement, with an overall adjusted Rand index of 0.936. We plotted the transmission proportion alongside the original paper’s results, finding that the estimations obtained with `sparrowhawk` are roughly similar per location. The different clustering implementation between `sparrowhawk` and the original article could explain part of these differences, in addition to different default parameters, and the lack of one assembly compared to the original study.

**Fig 4.**
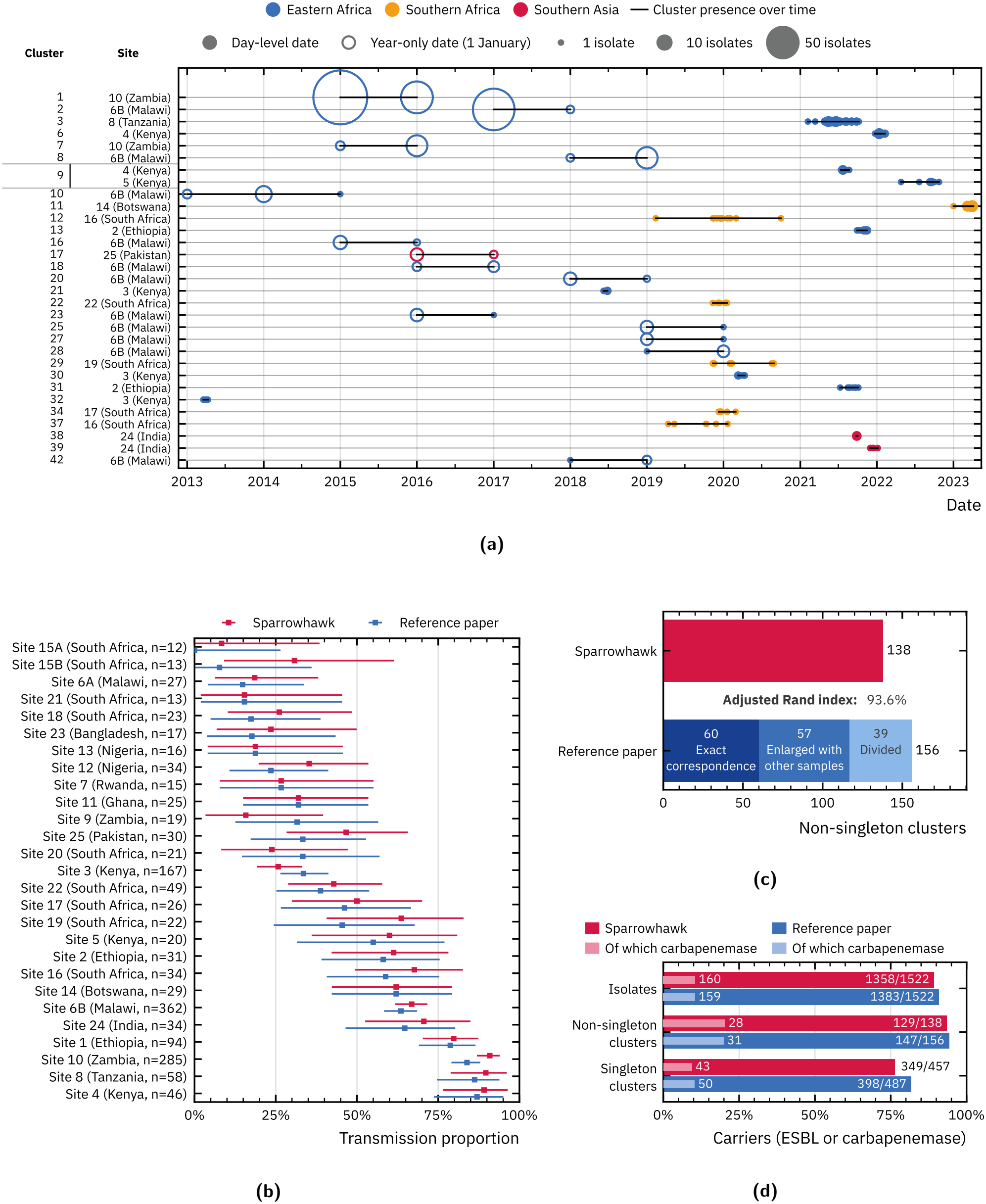
Comparison of results using `sparrowhawk` versus existing bioinformatics tools on transmission cluster identification and antimicrobial resistance gene detection. **a**, Cluster evolution with respect to date. Only the first thirty clusters, separated by location when their members are in multiple, and in which isolates appear in two or more different times are shown. X-axis dates indicate the start of each year. **b**, Transmission proportion of the different sites estimated from the clusters data, ordered by the reference paper values. This plot corresponds to fig. 1-B of [35]. **c**, Number of non-singleton clusters and adjusted Rand index. Reference-paper values are separated on those exactly reconstructed, enlarged with other samples, and those that are divided in various clusters. **d**, Number of isolates and clusters that have an extended-spectrum beta-lactamase or carbapenemase antimicrobial resistance (AMR) gene identified.

The study used Kleborate [38] to assess the AMR content of these genomes. To compare with this we developed the `sparrowhawk-amr` module: a lightweight reference gene detection method that allows search against a minimised reference database, which can be used to identify known resistance, virulence genes (but supports arbitrary targets). AMR detection methods rely on high-quality, expert-curated databases, such as AMRFinderPlus and the Reference Gene Catalog [39], ResFinder [2], or CARD [40]. Unfortunately, these are usually much larger than the standard assets one can ship with a website, and so their gold-standard detection methods cannot be directly adapted to run in a web browser. `sparrowhawk-amr` overcomes this by (offline) creating a reduced-size *k*-mer index built on the AMRFinderPlus database and with a total size of 15*≈* MB. Query sequences then have their nucleotide *k*-mers streamed (online) and exact matches are performed against the reference index, detecting indexed genes when the number of relative *k*-mer matches exceeds a preset threshold.

This tool is integrated into the “Gene calling” tab of `sparrowhawk`, where we implement a WebAssembly port of the `Prodigal` [41] algorithm for predicting protein-coding genes in bacterial genomes. After candidate genes are identified, we then scan their sequence content to match with the provided reference database.

Using AMRFinderPlus as the reference for ground truth, we evaluated `sparrowhawk-amr` (figure 5) on 10000 genomes, obtaining an F1 score for AMR gene detection of 92 %, with a precision of 91 % and a recall of 92 %. Including stress and virulence genes, the overall F1 score is 88 % (precision 91 %, recall 86 %). This shows that `sparrowhawk-amr` is able to provide most of the same AMR gene detection calls of AMRFinderPlus, while adjusting to the constraints of a website. These differences with the AMRFinderPlus references could be partly explained by high fractions of shared *k*-mers between gene alleles, or matches missed due to not allowing for small changes in the sequence. Further developments could explore the usage of split *k*-mers, or multiple *k* values to tackle these challenges. In the evaluation of fig. 4d, results show that `sparrowhawk-amr` is able to detect similar amounts of AMR genes related to extended-spectrum betalactamases (ESBL) or from carbapenemases to those from the original study.

**Fig 5.**
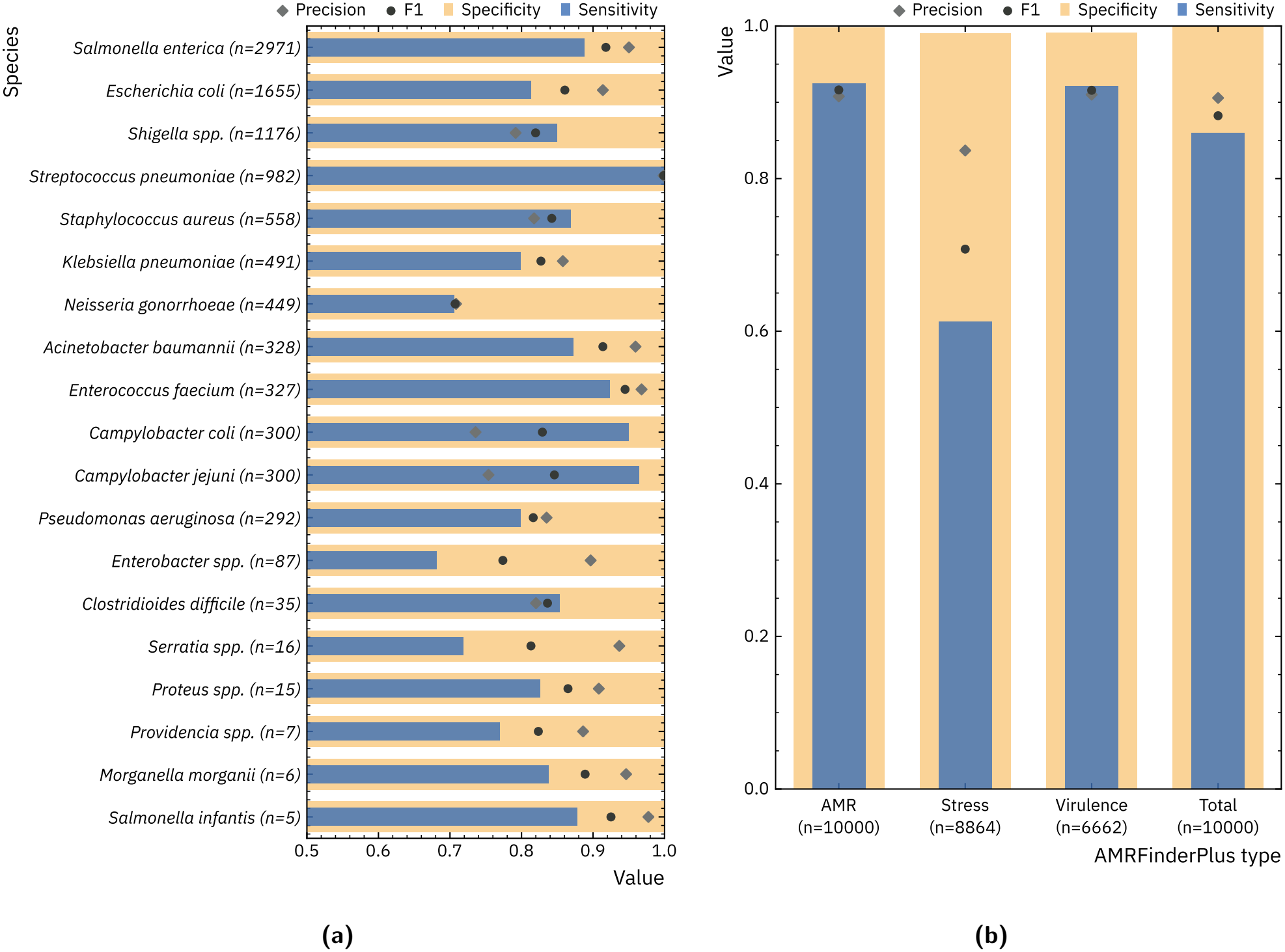
Precision, specificity, sensitivity (or recall), and the F1 score of `sparrowhawk-amr` evaluated on 10,000 real assembled genomes from public repositories against AMRFinderPlus as reference. **a**, Results per taxonomical groups (species and genus), with the number of samples per each one (*n*). **b**, Results per reference identified gene category: virulence, stress, or antimicrobial resistance, as well as the global evaluation. *n* shows here the number of samples that contain genes belonging to each group.

### Lowering the barrier for WebAssembly implementations of scientific methodologies

In `sparrowhawk` we show that it is possible to deliver a diverse set of high-performance methods as a web application. In addition to genome assembly, mapping and alignment, taxonomic identification, gene calling, and AMR detection, we have integrated other existing tools. One of them is `Deacon` [42], that allows filtering (or extracting) reads at scale and speed, particularly useful for e.g. removing human reads before analysis or data sharing, with the reads not leaving the device at any moment and thus preserving privacy. Another tab we included in `sparrowhawk` is “Embed proteins”, where users can do inference on a protein language model, ESM2 [43], and project the embeddings onto a two-dimensional space built with the Swiss-Prot dataset [44]. The inference can run on the CPU or can use the GPU thanks to the open standard WebGPU, speeding up inference. Furthermore, WebGPU is vendor-independent, so acceleration does not depend on a particular GPU, increasing the accessibility of this hardware acceleration. These tools both expand the functionality of `sparrowhawk` itself, and also demonstrate its adaptability as a general platform.

When adapting or creating a bioinformatic method for WebAssembly compilation, some common difficulties arose that required specialist attention to overcome. The main computational constraints are the maximum memory limitation per WebAssembly process (4 GB, recently grown to 16 GB in browsers that support 64-bit memory addresses), security of reading user input files, expected reduced performance against native executables (see tab. 1), complications when parallelising tasks, and the total amount of data one can comfortably ship with a website to avoid it being too heavy. For method developers in science, the adaptation of the code itself to successfully compile to WebAssembly, the compilation toolchain itself, and the interface with the web code (usually in a different language such as JavaScript) is not straightforward. This is compounded when GPU acceleration is necessary for more computationally intensive methods.

**Table 1.** Execution time performance comparison between devices (computer, smartphones), operating systems (GNU/Linux distribution, macOS, Windows), and Internet browsers (Mozilla Firefox, Chromium and Google Chrome, and Apple Safari), and also against native binaries. NA, unsupported; -, did not finish. Units are minutes:seconds.

|  | GNU/Linux |  |  | macOS |  |  |  | Windows |  | Android |  | iOS |
|---|---|---|---|---|---|---|---|---|---|---|---|---|
| Test | Native | Firefox | Chromium | Native | Firefox | Chrome | Safari | Firefox | Chrome | Firefox | Chrome | Safari |
| Assembly | 00:13 | 00:21 | 00:22 | 00:09 | 00:16 | 00:16 | 00:22 | 00:23 | 00:26 | 01:18 | - | 00:32 |
| A. (multi-*k*) | 01:56 | 02:52 | 03:13 | 01:37 | 02:23 | 02:31 | 02:42 | 03:17 | 03:50 | 12:37 | - | - |
| Embedding | 00:15 | 00:26 | 00:29 | 00:16 | 00:25 | 00:19 | 00:21 | 00:27 | 00:30 | 01:13 | 01:12 | 00:28 |
| E. (+GPU) | 00:01 | 00:12 | 00:03 | 00:05 | 00:10 | 00:05 | 00:13 | 00:23 | 00:09 | NA | 00:51 | NA |

We tackled all of these challenges in `sparrowhawk`, in a manner which we aimed to generalise. For example, we separate the method’s code from the website and compile and ship automatically multiple compilation targets when needed. When the user loads the website from a browser without support for one of them, the other is automatically used, extending usage. To further optimise memory usage, we distribute tasks across independent web workers. For example, in the “Alignment” tab, we effectively circumvent the memory limitation as it applies per process. The novel methods included have been designed for low memory footprint and execution times, as well as optimising the databases shipped where relevant (e.g. taxonomic identification). For applications where data security is paramount, `sparrowhawk` also provides an offline alternative which can be used without an internet connection (using Electron). This also opens the future possibility of distributing larger databases with the package.

To enable other method developers to reuse these techniques to port other tools, we have written a detailed development guide (https://sparrowhawk-docs.bacpop.org). This is a step-by-step document to create a simple bioinformatics method example from scratch, taking advantage of our integration approach.

As agentic programming became available during this project, we also recognised the importance of working with these tools directly, as they have enormous potential to further lower the barriers to advanced tool use. Our experience has been that these tools excel at multi-language integration required by WebAssembly projects. To formalise this, we have written a skill for coding assistants (available in the main code repository), which we refined through two different examples. First, the adaptation of another bioinformatic method: the read aligner `rammap`, based on `minimap2` [18, 19] (https://vrbouza.github.io/rammap-web). Second, and to showcase the interdisciplinary applications of `sparrowhawk`’s approach, a physics statistical analysis: CMS Combine [21] (https://vrbouza.github.io/combine-web). This skill has been used to successfully integrate `Mandrake` [20] into `sparrowhawk`, adding the functionality of fast embedding of large genome collections. The final optimised “rust-wasm-vue-tool” skill was able to create a correct WebAssembly port of `Mandrake` using a single prompt.

## Discussion

Computational analysis of biological data is too often restricted to research groups which specialise in this domain. Enabling easier access to the high-performance methods needed to analyse modern biological data through more visual tools (e.g. RStudio and Galaxy) has significantly broadened access. But even these approaches still require computational expertise to run, use, and interact with results. Creating web browser versions of computational tools is appealing due to their wide availability and familiarity, but is often not feasible with large or sensitive datasets, and adds additional development expertise and maintenance burden onto method developers.

In `sparrowhawk`, we have created a platform for performing computational analyses through a website using only local computation, without uploading data to remote servers. The compilation to WebAssembly of the different methods included allows our platform to run with only static server resource usage, in different Internet browsers, operating systems, and devices (from normal computers to tablets or smartphones). These features make WebAssembly a good solution for different contexts, such as data sovereignty concerns, analysis of sensitive data, or resource-constrained environments. Additionally, `sparrowhawk`’s interface is designed to be intuitive and simple, assisting those researchers not well-versed in bioinformatics or high performance computation. We also offer an Electron executable that can be downloaded for offline access to the tools provided.

To create `sparrowhawk` we have developed novel methodologies so that they could fit the constraints of running on a website. Among these we developed a fast and lightweight *de novo* bacterial genome assembler, and fast AMR gene signature detections using *k*-mers. We have also implemented sequence mapping and alignment, transmission clustering, taxonomic identification, gene calling, and read filtering for host depletion (with `Deacon`). We also provide an implementation of a large protein language model (ESM2) running through WebAssembly in a web browser, with GPU acceleration.

At the beginning of this work, it had already been acknowledged that WebAssembly had potential to remove barriers to computational method uptake [16]. But despite its many opportunities, WebAssembly poses challenges for developers, which has likely prevented its broader uptake. The adaptation of existing code for its compilation can be non-trivial, and developing a thoughtful graphical interface requires different programming skills. `sparrowhawk` lowers the barrier of introducing this technology in three main ways: by example, by our guide to offer a proven path on how to develop or adapt existing methodologies to work with WebAssembly on web browsers, and by providing instruction files to help develop these with large language models. These tools will be useful for people inside and outside bioinformatics, as we have shown by automatic porting of further bioinformatics and particle physics statistical methods. Through careful development of developer documentation and integration with agentic programming, we believe we have removed major constraints for creating web tools from command line applications. Running in the browser also has costs, for example CPU bound tasks were 1.2*−* 2.4*×* slower than native execution (tab. 1). The major remaining limitation of our approach is that it is centred around packages which can be implemented in the Rust programming language. However, this is not intrinsic to WebAssembly as the other popular scientific programming languages C, C++, R, Python, and Julia can all be compiled to this target. Additionally, agentic programming can be used for rapid language conversion, as shown by a recent effort which ported 40 popular bioinformatics tools to Rust [45].

We believe that the number of WebAssembly-designed methodologies will continue to grow in bioinformatics, and in many other scientific fields. The first beneficiaries will be the users of the case studies, for example genomic epidemiology or bacterial genomics researchers. The `sparrowhawk` template library and agentic skill will then help increase the impact of computational method developers’ research. Eventually, many more users across research, teaching, and public outreach stand to gain from the WebAssembly tools subsequently produced by these methodological researchers. Ultimately this work will better democratise computational methods in medical research, empowering those who generate data.

## Methods

### Short-read bacterial genome assembler: sparrowhawk-asm

`sparrowhawk-asm` is a de Bruijn graph assembler for short reads. By default it assembles iteratively over increasing values of *k* (multi-*k* mode); a single *k* can be chosen instead, through a command-line argument or a checkbox in the browser. Multi-*k* runs start at *k* = 21 and use a predefined set of five values between 21 and 71, extended up to a value close to the maximum read length. Single-*k* runs accept any odd *k* up to 255. The maximum read length and the range of base qualities are estimated from a sample of 5,000 reads. Each iteration has two stages: *k*-mer preprocessing (extraction, counting, and filtering), and graph construction and correction. The same Rust code is compiled natively and to WebAssembly; the differences between the two builds are summarised at the end of this section.

#### *k*-mer extraction and counting

As in `ska.rust` and `sketchlib.rust`, *k*-mers are extracted from each read using only bases with a Phred quality above 20 (the default quality floor), and never across ambiguous bases (`N`). Each *k*-mer is hashed with the `ntHash2` rolling hash [46], which also gives the hash of its reverse complement directly and is reused later to build the graph. Because we observed hash collisions at large *k* in large bacterial genomes (e.g. *Streptomyces avermitilis*), we increased the number of internal bit-wise splits per hash from two (the `ntHash2` default) to six, which substantially reduced hash collisions (see supp. mat. note 1).

In the same pass, we build a sketch of all *k*-mers, including those that fail the quality floor. Each sketched *k*-mer records the highest quality floor it passes, so that spectra at lower floors can be computed later without re-reading the data. In native runs, reads are also stored in memory using four bits per base: two for the nucleotide, and two encoding either an ambiguous base or one of three quality bins derived from the initial read sample. Because bacterial genomes are small (low coverage reads can be in the tens of megabytes, whereas high coverage experiments yield `fastq` of *≈*250*−* 1000 MB), this makes it feasible to hold the full read set in memory, even at high coverage.

*k*-mers are counted with a Bloom filter and a hash map, as in `ska.rust`. In native runs, these steps are parallelised: one thread reads the input in blocks, which are exchanged through two buffers with worker threads that hash and count, so that reading overlaps with hashing, the most CPU-intensive step.

#### Selecting the count threshold

*k*-mers are then filtered by count, using the *k*-mer spectrum (the histogram of *k*-mer counts) computed from the sketch to avoid Bloom filter false positives. Ideally, the spectrum consists of an error component, dominated by *k*-mers seen once, followed by a Poisson-distributed genomic peak at the mean *k*-mer coverage, 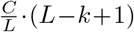 for average base coverage *C* and (constant) read length *L*, with smaller peaks at its multiples arising from repeats [47]. In real data, however, genomic peaks are usually overdispersed [48], under-sequenced GC-rich regions shift some genomic *k*-mers to lower counts [49], and the error component has a heavier tail than a Poisson distribution. Larger values of *k* lower the *k*-mer coverage, moving the genomic peak towards the error component, and make the spectrum sparser. The quality floor also shapes the spectrum: it splits reads into shorter fragments and so reduces the number of *k*-mers. Lowering it can separate the two components, but also fills the valley between them. Separating error from genomic *k*-mers is therefore difficult, particularly at large *k*. Other assemblers, such as `SPAdes`, `SKESA`, and `Minia`, rely mainly on their graph error correction logic, with `SPAdes` and `SKESA` also carrying per-*k*-mer quality information. Given our browser-imposed memory constraints, this approach is not feasible, nor is offloading data to disk, as `SPAdes` and `Minia` do.

For each *k*, `sparrowhawk-asm` therefore selects both a minimum count and a quality floor, combining model fitting with heuristics:

1. An empirical valley and genomic peak are located directly in the histogram, and accepted only if they pass checks on the mass and relative height of the peak.
2. Seeded with these values, four mixture models are fitted to the histogram by maximum likelihood using Nelder–Mead from 60 starting points [48, 50]. The error component has a free weight for the first bin and either a Pareto or a Weibull tail. The genomic component has two negative binomials, for the main peak and the first repeat peak, optionally with a modified beta-binomial that captures a heavier left shoulder and a stronger valley. The model is selected by the Bayesian information criterion (BIC) among those with plausible parameters. The fitted components are then used to move the empirical threshold to the left, so that at most one genomic *k*-mer is expected to be lost, provided the threshold stays above both the count at which the expected number of error *k*-mers equals the height of the genomic peak, and an absolute minimum of 2.
3. If no acceptable fit is found, if more than 5 % of the retained *k*-mers are expected to be errors, or if the genomic peak is below 10, the procedure is repeated at lower quality floors using the sketch, and heuristics select the best combination of threshold and floor. If a lower floor is chosen, *k*-mers are re-parsed at that floor and the threshold is refitted. Further fallbacks handle cases such as very low coverage, where the error and genomic components cannot be separated.

The retained *k*-mers are afterwards stored in a data structure designed for parallel neighbour search. This speeds up the graph construction step.

#### Graph construction and correction

For each retained *k*-mer, `sparrowhawk-asm` rolls its canonical and non-canonical hashes forwards and backwards over the four possible bases to find its neighbours. It then builds a node-centric de Bruijn graph [51, 52], with one node per *k*-mer carrying its count and edges between overlapping *k*-mers, using `sparrowhawk-graph`, a Rust crate we developed to modularise the graph data structures and common functions related to de Bruijn graphs.

The assembly graph is then corrected with approaches adapted from `SPAdes` and `Minia`. Dead ends [31, 53], short branches without continuation which are common at read ends, are removed when they are very short or their coverage differs strongly from that of their surroundings (with thresholds that scale with *k*). Bubbles and bulges, where paths diverge and rejoin, are collapsed when the mean *k*-mer counts of their branches differ substantially, as expected from residual sequencing errors rather than repeats. Erroneous connections, low-coverage links between two regions of much higher coverage (four-fold by default), are removed. These corrections are repeated, interleaved with merging non-branching paths into single nodes to reduce the size of the graph, until no further dead ends or bubbles can be removed. The corrected graph is written in DOT, GFA1, and GFA2 formats.

#### Contig extraction and multi-*k* iteration

Contigs are the non-branching paths that remain after removing junction *k*-mers. In single-*k* mode, contigs shorter than 500 bp are discarded, and the sequences of the rest are reconstructed from the stored *k*-mer information and written as `fasta`. In multi-*k* mode, all contigs are kept in a compact 2-bit encoding, and their *k*-mers are added to those from the reads in the next iteration, carrying forward the structure resolved at smaller *k*. Later iterations also inherit preprocessing decisions: if two consecutive iterations lower the quality floor, the lower floor becomes the default. Iteration stops early in two cases. If no threshold can be found at any quality floor, the assembly is still attempted, but the previous iteration’s contigs are returned unless the empirical area under the N curve (auN) increases. Iteration also stops if the fitted or empirical genomic peak falls below four.

#### WebAssembly build

`sparrowhawk-asm` compiles to the `wasm32-unknown-unknown` and `wasm64-unknown-unknown` targets with minimal changes: an interface *struct*, `AssemblyHelper`, communicates with the website, and a few dependencies needed small modifications to compile. The browser build differs from the native one in two aspects. To better accommodate to the web constraints, reads are not kept in memory, so the input is re-read if the quality floor is lowered. Secondly, all runs are single-threaded, which affects *k*-mer counting, the model fits, and neighbour search.

#### Benchmark

We evaluated `sparrowhawk-asm` on real and simulated reads from six species: *Escherichia coli, Mycobacterium tuberculosis, Neisseria meningitidis, Staphylococcus aureus, Streptococcus pneumoniae*, and *Streptomyces coelicolor*. These include widely studied species and priority pathogens [54], as well as difficult to assemble genomes with many repeats such as *N. meningitidis* [55], and one with a linear chromosome of 8.7 Mbp (*S. coelicolor* [56]). For real data, we sampled 30 paired-end read sets with 150 bp reads per species from ENA [57] (180 in total), excluding those with adapter or primer sequences or low quality according to `fastp` [58]. To assess performance at low coverage, three read sets per species were chosen at random and sub-sampled to 5*×*, 10*×*, 15 *×*, and 20*×* coverage. For simulations, we generated 30 read sets per species from one reference genome with each of `ART` [59] and `Mason` [60]. All read sets were assembled with `sparrowhawk-asm`, `SPAdes` [28], `SKESA` [29], `Minia` [30], and `Velvet` [31] and evaluated with `QUAST` [61], using the reference genomes for the simulated data. We evaluated contiguity using the *Nx* function (*N* (*x*)), which is defined as the length for which the collection of all contigs of that length or larger covers at least the *x* % of the whole assembly. Detailed results of the benchmark can be also seen in supp. figs. 1–18, and the real read sets and reference assemblies in supp. tabs. 1 and 2.

### Aligning and mapping within species

We adapted `ska.rust` for `sparrowhawk` using two small object wrappers that represent the data needed for mapping and alignment respectively. To allow a larger number of samples to be processed for the alignment tab of the website, we optimised the inner data storage and implemented parallelisation using web workers. Additionally, we have included an alignment viewer based on `MSEABOARD` [36], that allows users to visualise the read mapping result. In the “Alignment” tab, phylogenetic tree construction is provided based on the alignment that `ska.rust` produces. We first calculate a triangular matrix of distances as a flattened vector between the given samples (assembled genomes, or reads) based on the SNP differences from the alignment. Then, we use a distance-based algorithm to reconstruct a phylogenetic tree through a neighbour-joining approach. The resulting tree is visualised in the web platform using `Taxonium` [37].

### Transmission clustering

The transmission approach infers possible clusters simply by clustering isolates with closely related SNP counts across samples. Also known as single linkage clustering, a graph with nodes representing samples, and edges added whenever the distance between samples is below a user-specified threshold. The connected components are the possible transmission clusters. If optional metadata (ID, date, and location) is provided, `sparrowhawk` is able to improve the presentation of results, as shown in supp. fig. 31. It is possible to run the transmission clustering workflow both in the “Alignment” and “Transmission” tabs of the website.

The data for the replication study of [35] was downloaded from ENA. Of the 1523 genomes reported on the article, 1522 were available in public repositories (either as reads or assemblies) and used for the comparison (supp. mat. tab. 3). The temporal position was set extracting it from their ENA accessions, although some had only year-level resolution (and got as date the first of January of the reported year).

### Inverted indexing for taxonomic identification

We constructed a large inverted index with `sketchlib.rust` using high-quality assembled bacterial genomes from the AllTheBacteria (`AtB`) [34] dataset, clustered by sequence similarity using `Gemsparcl` [33]. This yielded 7305 clusters, each with its own identification number. To select a representative from each cluster, we ran `QUAST` and selected the assembly with the best area under the *Nx* function value (see above). With those 7305 cluster representative assemblies, we used `sketchlib.rust` to create an inverted index, with each entry labelled with metadata representing the species name. For each cluster, we incorporated the list of the Genome Taxonomy Database (GTDB) [62, 63] species assignment using `GTDB-Tk` [64, 65] of each cluster’s assemblies, how many assemblies of each taxonomical assignment there were, and the `Gemsparcl` cluster identification number.

Online, when query files (either `fasta` or `fastq`) are provided, they are first sketched and the result is compared against the inverted index. The three leading results are provided, along with the metadata and the average nucleotide identity (ANI) of the match. If there is a tie in any of the three ranks, all the clusters with the same level of agreement are reported as well.

The implementation returns a simplified taxonomic assignment per result, which can be a species, a genus, or “Unknown”. Heuristically, we found that if there are more than 14 taxonomical assignments, the cluster usually belongs to one unique species, or many very small components (less than *≈* 0.1 %), so we take the leading component as the simplified taxonomic assignment. Before reporting it, we first modified by removing the derivations defined with underscores that exist on GTDB. In clusters with less than 14 assemblies, we remove all GTDB suffixes (e.g. A) as well and deduplicate the resulting list. If only one element survives, that is the reported species that can also be “Unknown” if that is the assignment. If not, we see if the genera of the assignments are the same: if it is, we report in this field “*Genus spp*.”. If not, we report “Unknown”.

We evaluated parameter selection for the inverted index, the length of the *k*-mers used (*k*) and the sketch size *ss*, using the whole `AtB` dataset (2339195 assemblies, release 0.2) as evaluation data. Our aim was to optimise for the best accuracy, with an inverted index that was small enough to fit as a website asset (maximum *≈* 10 20 *−*MB). We found *k* = 17 and *ss* = 50 to be a reasonable compromise, providing an inverted index of 3.49 MB and an accuracy of 97.8 %.

### Adapting external tools for WebAssembly

To provide fast read filtering, we adapted `Deacon` [42] for compilation to WebAssembly. A bridge package that imports this library was built for its integration into `sparrowhawk`. The web interface allows to filter (and remove), or select reads from a larger set. To do so, usually large indexes are used (10^2^ *−*10^3^ MB), that are not suitable to directly be shipped with a website. Therefore, we ask the user to drag-and-drop a predownloaded index first, followed by the read file they wish to decontaminate.

`sparrowhawk` includes gene calling through the `Prodigal` [41] algorithm, implemented with an existing Rust port (`orphos` [66]) that includes performance improvements. We followed `Deacon`’s integration approach and created a bridge package that contained the final interface to connect it to the web code. The results can be retrieved and downloaded as `gffs`, and we have also included a genome viewer based on `JBrowse2` [67] to visualise the gene calls.

### Antimicrobial resistance detection with sparrowhawk-amr

`sparrowhawk-amr`’s algorithm is based on an index built from *k*-mers extracted from genes of a reference database. In our case, we use AMRFinderPlus [39] databases (for the `sparrowhawk` release here depicted, we used the database version 2026-05-15.1), allowing for convenient updates. Although other options such as protein *k*-mers were explored, the final integration on `sparrowhawk` uses nucleotide sequences from genes with *k* = 31. The index is constructed from the database by extracting the coding sequences (CDSs) of the AMR genes, as well as other non-AMR genes included (stress resistance, and virulence), but not considering point mutations, either in CDSs or inter-genic regions (IGRs). *k*-mers are then streamed from each gene’s CDS and stored temporarily in memory. `sparrowhawk-amr` selects “report units” in the index. They are not necessarily the exact genes from the AMRFinderPlus database, as many *k*-mers might be shared across genes, making thus resolving those differences hard. We identify those *k*-mers that are unique per gene, and only include genes that have more than a particular threshold (21 unique *k*-mers by default). In the case a gene does not have that number of unique *k*-mers, we instead search for unique *k*-mers shared by all descendants of the most recent ancestor that contains class matches in AMRFinderPlus’s gene hierarchy. If those matches are larger than 30, we consider as reportable node also that hierarchy node. Both an intermediate node, and a tip (a single gene or allele) can happen to be reportable nodes. The index is constructed with the reportable entries, alongside metadata from AMRFinderPlus’s database, and finally serialised and compressed. `sparrowhawk-amr` is implemented in `sparrowhawk` for detecting AMR gene content in any DNA sequence, as well as being integrated with the gene calling feature where its results appear also as annotations in the genome viewer. Online, when a query sequence –a called gene, or an arbitrary sequence– needs to be assessed, we stream its *k*-mers and check the matches with the index. If the fraction of matches of a particular report unit over its total number is higher than a threshold (0.30 by default for exact gene alleles, and 0.10 for gene groups), we call that particular report unit.

`sparrowhawk-amr`’s evaluation was carried out using data from EMBL-EBI’s AMR portal [68] which contains experimentally measured AMR signatures. We sampled 10000 entries (supp. mat. tab. 4), ensuring representation of pathogens of interest (ESKAPEE: *Enterococcus faecium, Staphylococcus aureus, Klebsiella pneumoniae, Acinetobacter baumannii, Pseudomonas aeruginosa, Enterobacter spp*., and *E. coli*), and otherwise as much diversity of antibiotic classes as possible. We compared `sparrowhawk-amr` calls with those of AMRFinderPlus version 4.2.7 with database version 2026-05-15.1, which was also used for creating `sparrowhawk-amr`’s indexes. Performance was assessed by evaluating specificity, sensitivity, precision, and the F1 score.

### Embedding proteins with ESM2

We use a translated ESM2 [43] model with 8M parameters to the ONNX [69] format from HuggingFace (`Xue-Jun/esm2-8M`), whose architecture we translate to Rust-defined code with the `burn` crate, and whose weights are stored as `bpk` (burnpack). We checked the difference in embeddings (with 20 proteins) and in the weights of the model against the reference baseline, which we found to be negligible. The final model architecture file is included in the WebAssembly binary, while the model weights are shipped with the website, and occupy 13.7 MB (compressed).

To create the 2D projection of embeddings for visualisation, we use the `umap-learn` module from Python, and the Swiss-Prot dataset [44], composed of 575503 manually-curated proteins. To do so, first we embed each protein, truncated at a maximum of 1022 residues. Then, we train an UMAP on Python with the embeddings. The resulting reference projection is stored as a binary file that is shipped with the website. The user can choose to run the model for inference on their own data with CPU or GPU (using WebGPU if available). Resulting protein embeddings are projected into the previously trained UMAP space, yielding an interactive plot.

### Integration, web interface, and performance evaluation

The website components are built using JavaScript/TypeScript with the Vue framework. When building and running the website, all Rust methods are first compiled to WebAssembly automatically via `wasm-bindgen`, and afterwards the rest of the website is built. We also offer a traditional application packaging approach which allows full offline use of `sparrowhawk`. To do so, we make the web portable using Electron, which embeds the web browser Chromium. This process is automated in a single build command. Parallelisation is available for the “Alignment”, “Taxonomic ID”, and “Gene calling” tabs, by creating multiple JavaScript web workers, whose multiplicity can be set up with a slider. In these cases, multiple input files can be submitted at once.

We aimed to make the design of the website simple and intuitive, and generalisable across methods, to ease reuse by other developers. We offer a large space for the method selected and its output, which generalises well across tools. Though usage varies slightly across the tools, we found a common set of steps of asking the user to provide input data, running the method, and showing result visualisations, and finally saving the outputs to disk from memory. The main content panel is divided in a parameter vertical bar at the left, in which settings for each method are set before running, while at the right the user can drag-and-drop the input data. A small help box in the top left explains briefly the method, how to use it, and provides links to example input data that can be downloaded and tested. Other general questions are addressed by the frequently asked questions (FAQ) page, with a link in the bottom left corner of the methods bar. The design of the website has also been adapted for smartphones, where the contents are rearranged for better user experience. Screenshots of all tabs of `sparrowhawk` are shown in supp. figs. 19–30.

Computational benchmarking of `sparrowhawk` was done across different operating systems (OSs), and Internet browsers, running the two tasks that were more easily comparable between native and on browser execution. We used a set of *S. pneumoniae* read simulations for the assembly tasks, and 50 protein sequences for the embedding ones. GNU/Linux-based system checks and Windows-based checks used the same system: 64 GB of DDR5 RAM, an AMD Ryzen 7 9800X3D, and an AMD Radeon RX 6800 XT GPU with 16 GB VRAM. GNU/Linux distribution was Arch Linux, and the Windows version was Microsoft Windows 10. Mac-based system checks used macOS Tahoe 26.6, 24 GB RAM, and an Apple M5 Pro with 15 CPU threads and 16 GPU threads. Android checks used a Google Pixel 8 running GrapheneOS over Android 17, build 2026102101, and iOS used an Apple iPhone Air.

The results, on tab. 1, show a clear increase in execution time when moving from native to browser in both GNU/Linux and macOS, with browser execution being between 1.2 *×*and 2.4 *×*slower. This difference is particularly large when running the task with GPU acceleration, where the ratios reach 12 *×*. Between the GNU/Linux and Microsoft Windows systems, where the hardware is the same, browser execution times were faster in all cases on GNU/Linux. Between Firefox and Chromium-based browsers, available in all systems (on iOS all browsers use Safari’s WebKit engine, so only Safari was tested), results are mixed. Firefox was faster than or equal to Chromium-based for the assembly tasks and embedding on the CPU (this last except on macOS). When taking advantage of GPU acceleration with WebGPU, results inverted and Chromium-based browsers showed better performance, with Firefox being up to four times slower on GNU/Linux. This suggests a more efficient WebGPU implementation on the Chromium-based browser family.

On smartphones, results are harder to compare. Firefox was able to finish all tasks on Android except embedding with GPU acceleration, where we were not able to enable WebGPU support, even on the nightly version. Chrome was extremely slow on Android and did not finish the assembly tasks, while being able to do both the embedding with and without GPU acceleration. On iOS, Safari was able to run all tasks except embedding with GPU acceleration, as it did not support WebGPU. However, the multi-*k* assembly task failed because Safari does not support 64-bit memory addresses for WebAssembly, and the execution reached the memory limit.

## Supporting information

Supp. AMR evaluation assemblies table

Supp. assembler evaluation reads table

Supp. assembler evaluation references table

Supp. transmission samples table

## Acknowledgements

We thank Andrea Epifani from Gomoku Studio for creating the design of the web front end and improving its early functionality.

## Declarations

## Funding

This work was supported by the European Molecular Biology Laboratory’s European Bioinformatics Institute (EMBL-EBI).

## Conflicts of interest

No competing interest is declared.

## Data availability

Assembler benchmark accessions are available at supp. tabs. 1 and 2, as well as in the corresponding attached supplementary `TSV` files. Data used on the reproduced transmission clustering analysis is summarised on supp. tab. 3, and in the corresponding attached `TSV` file. Data used on the `sparrowhawk-amr` evaluation is summarised on supp. tab. 4, and in the corresponding attached `TSV` file. The AllTheBacteria dataset version used was 0.2 and it is available on its website, https://allthebacteria.org. Swiss-Prot version used was the release 2026 04, and it is available for download at UniProt’s website https://www.uniprot.org/help/downloads. Indexes used by `sparrowhawk` are shipped with it, and thus available with its code (see below).

## Code availability

`sparrowhawk-asm`’s code is available at https://github.com/bacpop/sparrowhawk-asm (`master` branch). `sparrowhawk-graph`’s code can be found in https://github.com/bacpop/sparrowhawk-graph (branch: `master`); `ska.rust`’s in https://github.com/bacpop/ska.rust (branch: `sparrowhawk-dev`), containing also the code for the transmission clustering application; `sketchlib.rust`’s in https://github.com/bacpop/sketchlib.rust (branch: `sparrowhawk-dev`); `sparrowhawk-amr`’s in https://github.com/bacpop/sparrowhawk-amr (branch: `master`). The `Deacon` code used can be found in https://github.com/bede/deacon (branch: `wasm1`). `orphos`’s code used is in https://github.com/vrbouza/orphos (branch: `victordev`). The web frontend, the Electron program, the documentation files (including the development guide), the agentic skill, and the integration of the different methodologies are located in the main repository of `sparrowhawk`, https://github.com/bacpop/sparrowhawk (branch: `main`). The code for the particle physics analysis example developed using `sparrowhawk` as reference can be found at https://github.com/vrbouza/combine-web (branch: `master`), and for the read aligner, in https://github.com/vrbouza/rammap-web (branch: `main`).

### Large language models (LLMs) usage

We used LLMs to draft Rust tests in various crates, perform code reformulations (e.g. moving existing handwritten code to a separate module or crate), and for checking the existence of potential undiscovered bugs. Also, to outline initial code following the authors’ architecture and guidelines in some subprojects, such as `sparrowhawk-amr`, protein embeddings, the Electron program, or other improvements done to particularly `sparrowhawk-asm` and `ska.rust`, as well as develop the examples of additional bioinformatics and physics methodologies. The final code in all cases was manually reviewed and improved afterwards by the authors. Different models were used (depending on the author and moment in time). Models: `Opus`, in its major versions 4 and 5 (Anthropic), and `GPT`, major versions 5 and 6 (OpenAI).

## Author contribution

Conceptualisation: J.A.L. Data curation: V.R.B. Formal analysis: V.R.B. Funding acquisition: J.A.L. Investigation: J.A.L., V.R.B. Methodology: J.A.L., V.R.B., A.A., J.H.v.W., B.C., E.E.O., L.N., V.G. Project administration: J.A.L. Resources: J.A.L., J.H.v.W. Software: J.A.L., V.R.B., A.A., J.H.v.W., B.C., E.E.O., L.N., V.G. Supervision: J.A.L. Validation: V.R.B., B.C. Visualisation: V.R.B. Writing (original draft): V.R.B. Writing (review and editing): J.A.L., V.R.B., A.A., J.H.v.W., B.C., E.E.O., L.N., V.G.

## Notes

### Competing Interest Statement

The authors have declared no competing interest.

