## Supplementary material for "Sparrowhawk: harnessing WebAssembly for private microbial genomics, and beyond"

### Contents

**Supplementary figures** **2**
sparrowhawk-asm benchmark . . . . . . . . . . 3

**Supplementary tables** **33**

**Supplementary notes** **36**

**Supplementary note 1: $k$-mer hash collisions in sparrowhawk-asm** **36**

### Supplementary figures

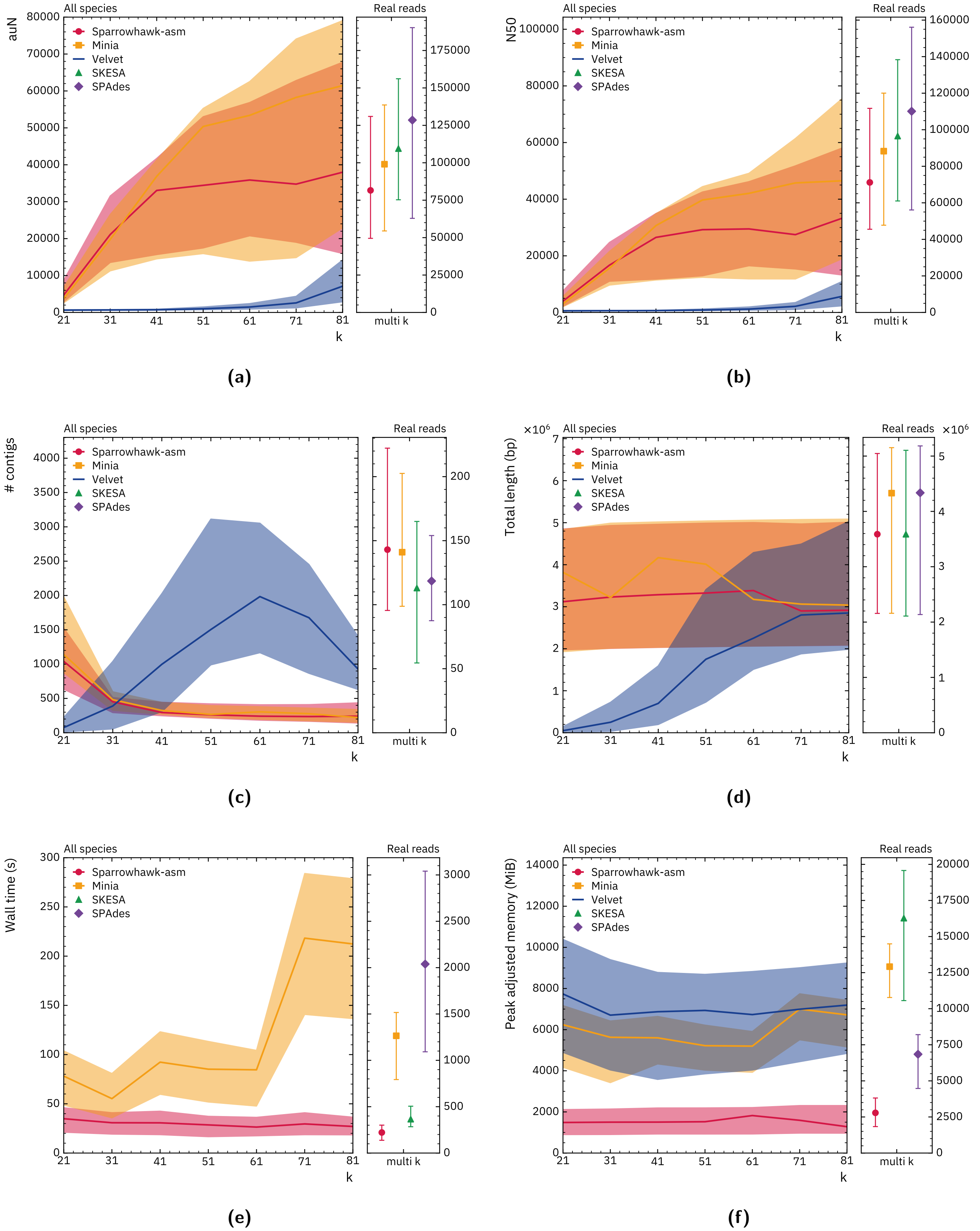


**Supplementary fig. 1.** `sparrowhawk-asm` benchmark on real reads, pooled over the six species (180 read sets): **a,** auN; **b,** N50; **c,** Number of contigs; **d,** Total length; **e,** Wall time; **f,** Peak adjusted memory. Lines show the median over read sets at each $k$-mer length, and shaded bands the interquartile range. The right-hand panel shows the multi-$k$ assemblers (`sparrowhawk-asm`, `Minia`, `SKESA`, and `SPAdes`) as medians with interquartile whiskers, on their own $y$ axis. The main $y$ axis is limited to 80,000 in **a** and to 300 s in **e**, where `Velvet` is also left out, to show the differences between `sparrowhawk-asm` and `Minia`. Peak adjusted memory includes both RAM as well as the file disk allocation. We use for this the peak memory charged to the run's control group (cgroup), which includes the in-memory (tmpfs) working directory used by `Minia` and `SPAdes`.

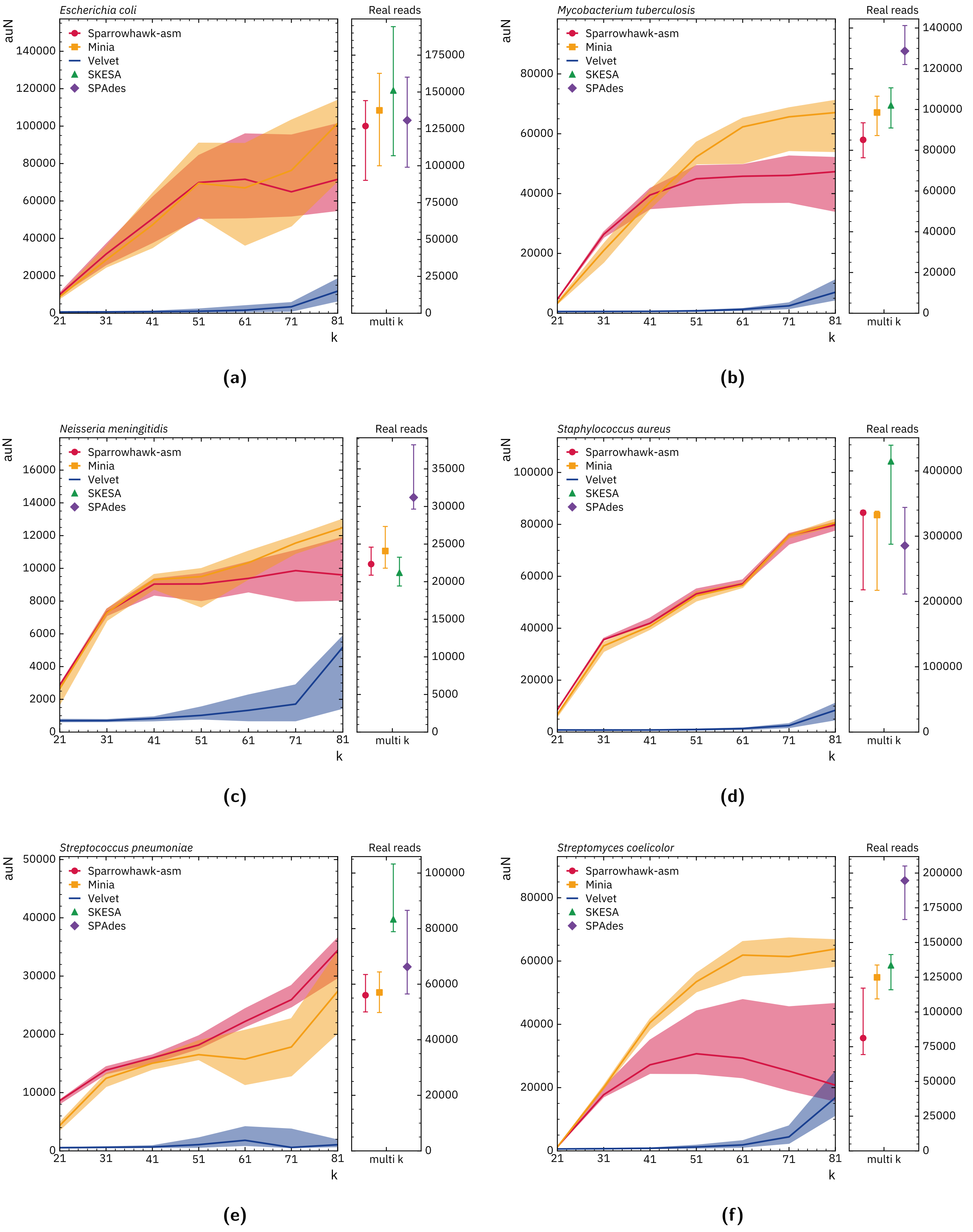


**Supplementary fig. 2.** `sparrowhawk-asm` benchmark on real reads: auN per species (30 read sets each). Lines show the median over read sets at each $k$-mer length, and shaded bands the interquartile range. The right-hand panel shows the multi-$k$ assemblers (`sparrowhawk-asm`, `Minia`, `SKESA`, and `SPAdes`) as medians with interquartile whiskers, on their own $y$ axis. **a,** *Escherichia coli*; **b,** *Mycobacterium tuberculosis*; **c,** *Neisseria meningitidis*; **d,** *Staphylococcus aureus*; **e,** *Streptococcus pneumoniae*; **f,** *Streptomyces coelicolor*.

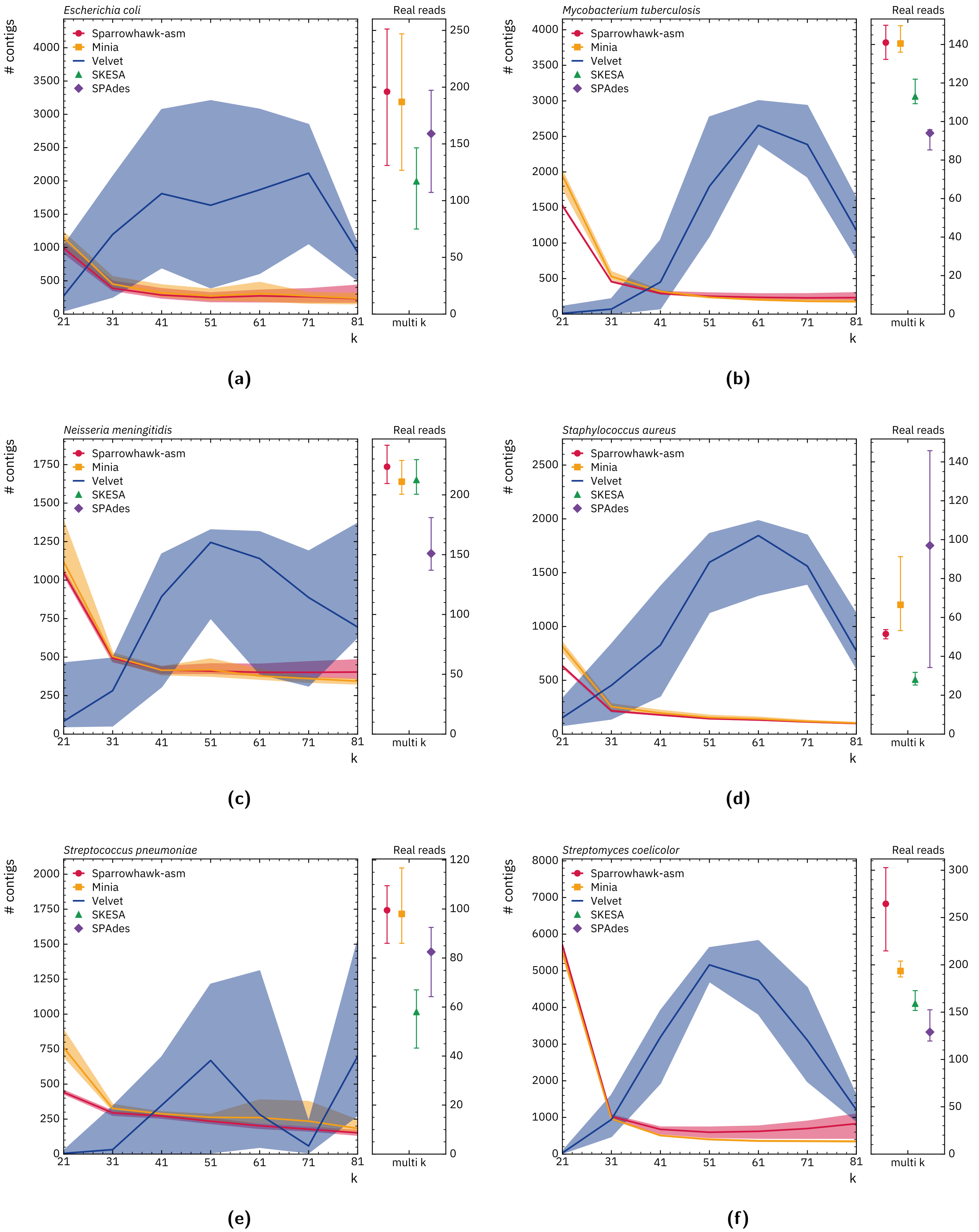


**Supplementary fig. 3.** `sparrowhawk-asm` benchmark on real reads: number of contigs per species (30 read sets each). Lines show the median over read sets at each $k$-mer length, and shaded bands the interquartile range. The right-hand panel shows the multi-$k$ assemblers (`sparrowhawk-asm`, `Minia`, `SKESA`, and `SPAdes`) as medians with interquartile whiskers, on their own $y$ axis. **a,** *Escherichia coli*; **b,** *Mycobacterium tuberculosis*; **c,** *Neisseria meningitidis*; **d,** *Staphylococcus aureus*; **e,** *Streptococcus pneumoniae*; **f,** *Streptomyces coelicolor*.

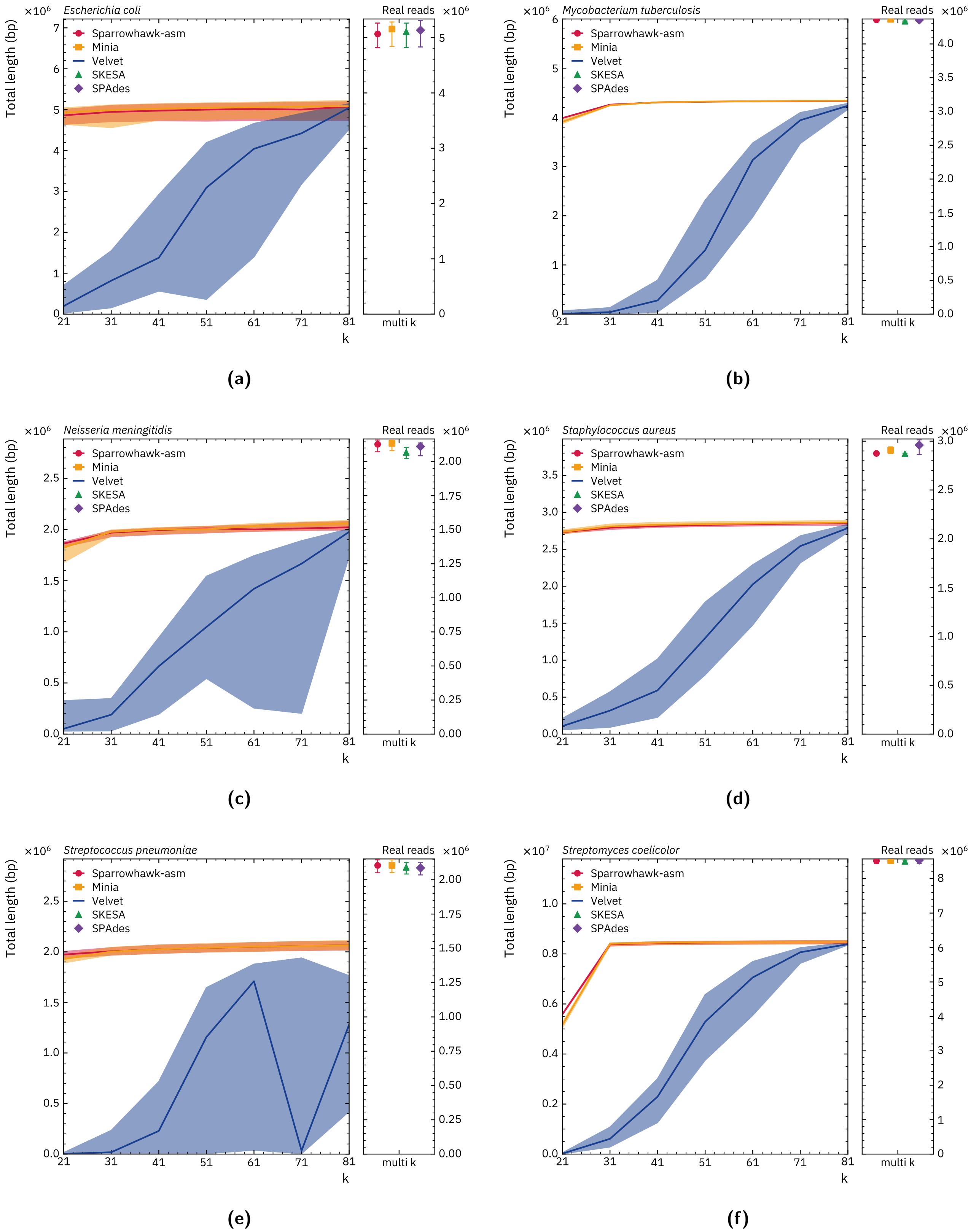


**Supplementary fig. 4.** `sparrowhawk-asm` benchmark on real reads: total assembly length per species (30 read sets each). Lines show the median over read sets at each $k$-mer length, and shaded bands the interquartile range. The right-hand panel shows the multi-$k$ assemblers (`sparrowhawk-asm`, `Minia`, `SKESA`, and `SPAdes`) as medians with interquartile whiskers, on their own $y$ axis. **a,** *Escherichia coli*; **b,** *Mycobacterium tuberculosis*; **c,** *Neisseria meningitidis*; **d,** *Staphylococcus aureus*; **e,** *Streptococcus pneumoniae*; **f,** *Streptomyces coelicolor*.

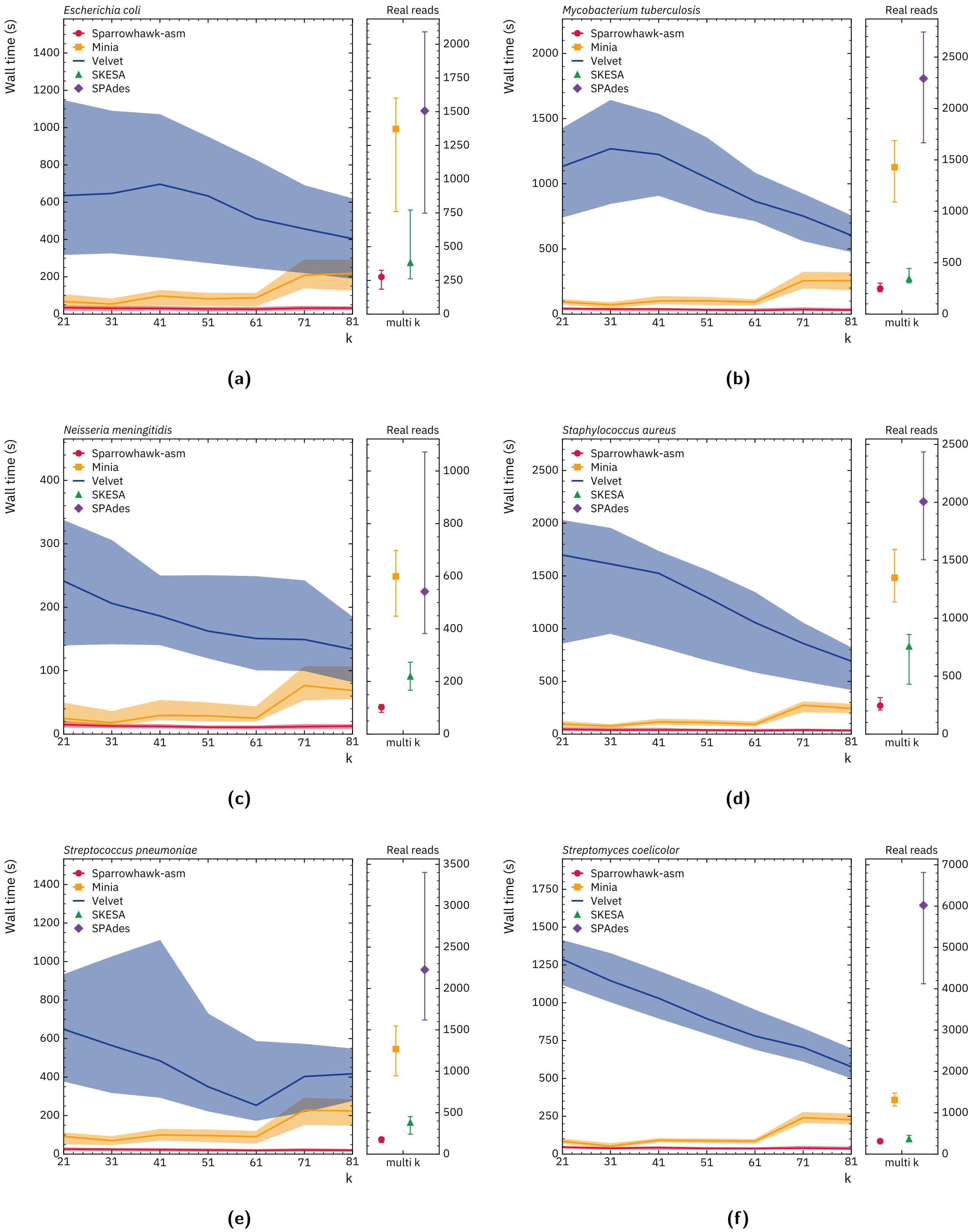


**Supplementary fig. 5.** `sparrowhawk-asm` benchmark on real reads: wall time per species (30 read sets each). Lines show the median over read sets at each $k$-mer length, and shaded bands the interquartile range. The right-hand panel shows the multi-$k$ assemblers (`sparrowhawk-asm`, `Minia`, `SKESA`, and `SPAdes`) as medians with interquartile whiskers, on their own $y$ axis. **a,** *Escherichia coli*; **b,** *Mycobacterium tuberculosis*; **c,** *Neisseria meningitidis*; **d,** *Staphylococcus aureus*; **e,** *Streptococcus pneumoniae*; **f,** *Streptomyces coelicolor*.

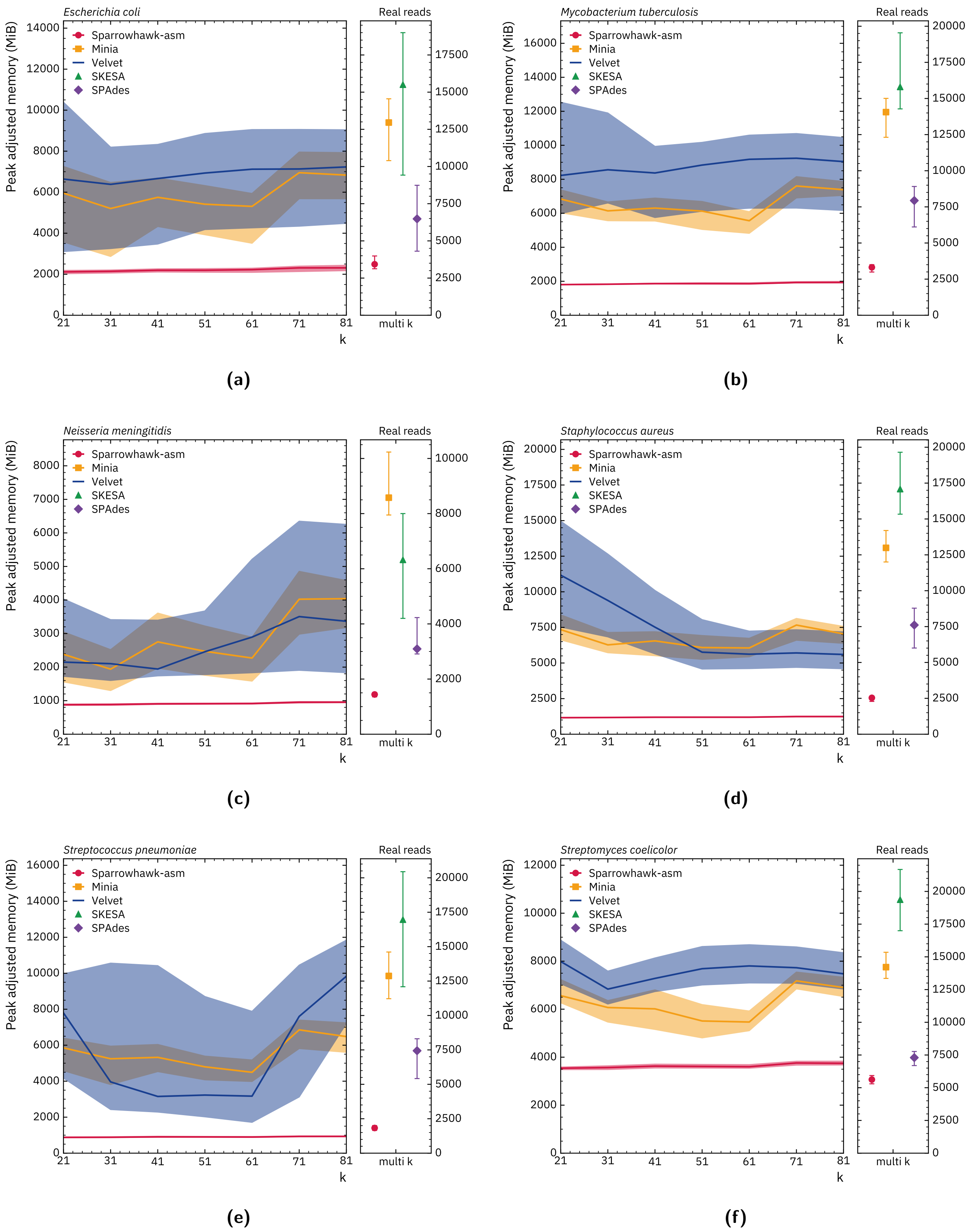


**Supplementary fig. 6.** `sparrowhawk-asm` benchmark on real reads: peak adjusted memory per species (30 read sets each). Lines show the median over read sets at each $k$-mer length, and shaded bands the interquartile range. The right-hand panel shows the multi-$k$ assemblers (`sparrowhawk-asm`, `Minia`, `SKESA`, and `SPAdes`) as medians with interquartile whiskers, on their own $y$ axis. Peak adjusted memory includes both RAM as well as the file disk allocation. We use for this the peak memory charged to the run's control group (cgroup), which includes the in-memory (tmpfs) working directory used by `Minia` and `SPAdes`. **a,** *Escherichia coli*; **b,** *Mycobacterium tuberculosis*; **c,** *Neisseria meningitidis*; **d,** *Staphylococcus aureus*; **e,** *Streptococcus pneumoniae*; **f,** *Streptomyces coelicolor*.

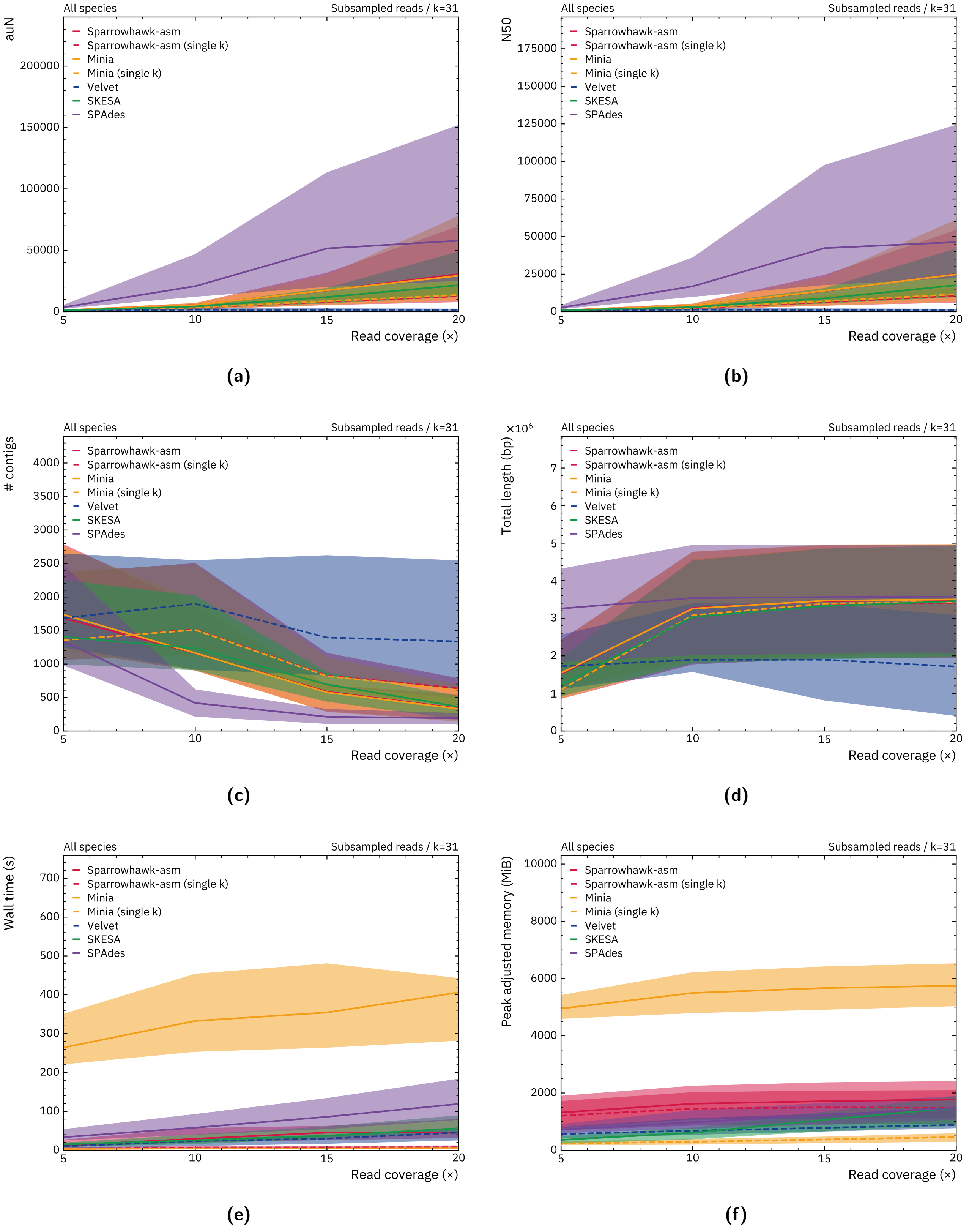


**Supplementary fig. 7.** `sparrowhawk-asm` benchmark on real reads subsampled to 5, 10, 15, and 20× coverage, pooled over the six species (three read sets per species and coverage): **a,** auN; **b,** N50; **c,** Number of contigs; **d,** Total length; **e,** Wall time; **f,** Peak adjusted memory. Lines show the median over read sets at each subsampled coverage, and shaded bands the interquartile range. Single-$k$ assemblers ran at $k = 31$. Peak adjusted memory includes both RAM as well as the file disk allocation. We use for this the peak memory charged to the run's control group (cgroup), which includes the in-memory (tmpfs) working directory used by `Minia` and `SPAdes`.

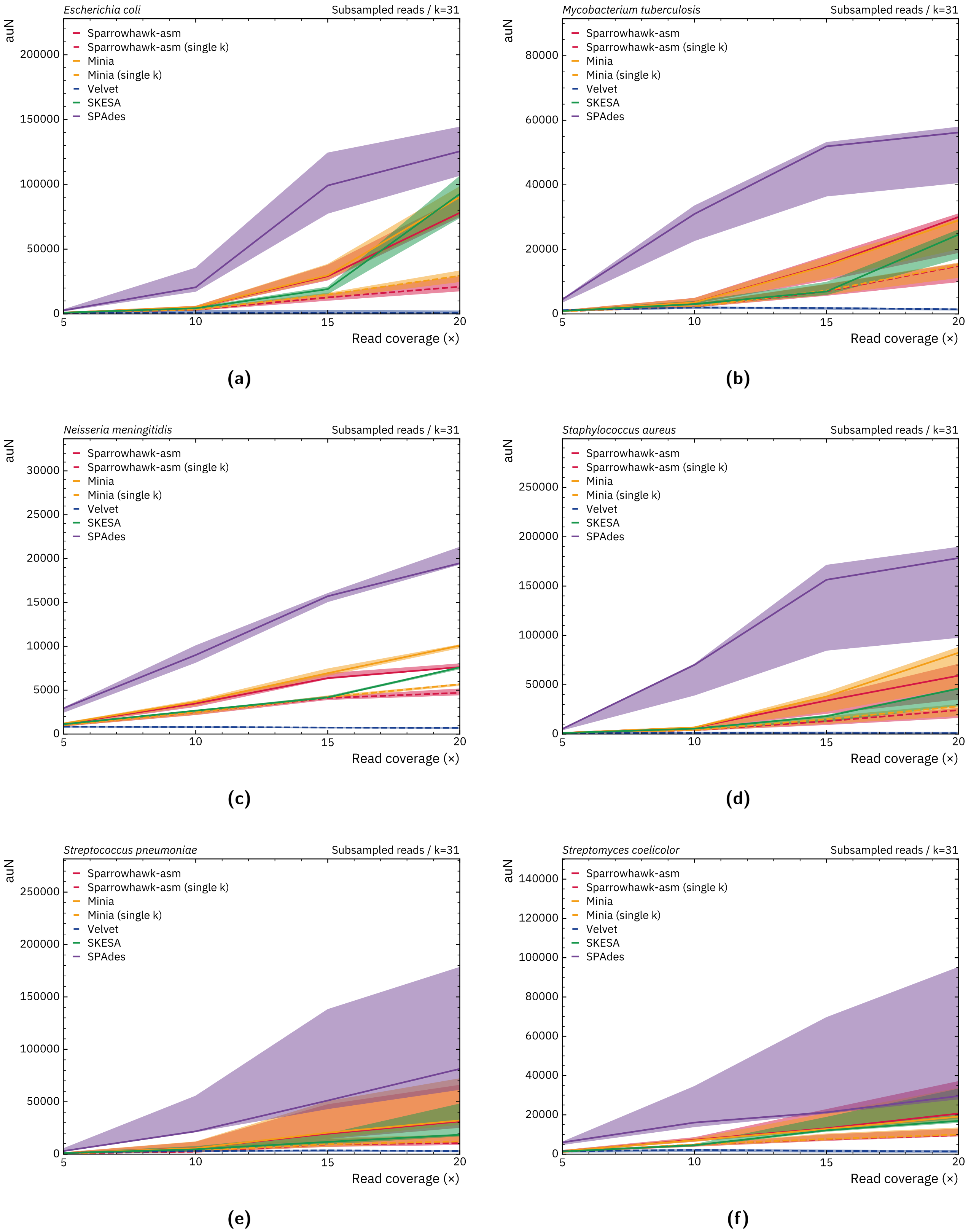


**Supplementary fig. 8.** `sparrowhawk-asm` benchmark on subsampled real reads: auN per species against coverage (three read sets each). Lines show the median over read sets at each subsampled coverage, and shaded bands the interquartile range. Single-$k$ assemblers ran at $k = 31$. **a,** *Escherichia coli*; **b,** *Mycobacterium tuberculosis*; **c,** *Neisseria meningitidis*; **d,** *Staphylococcus aureus*; **e,** *Streptococcus pneumoniae*; **f,** *Streptomyces coelicolor*.

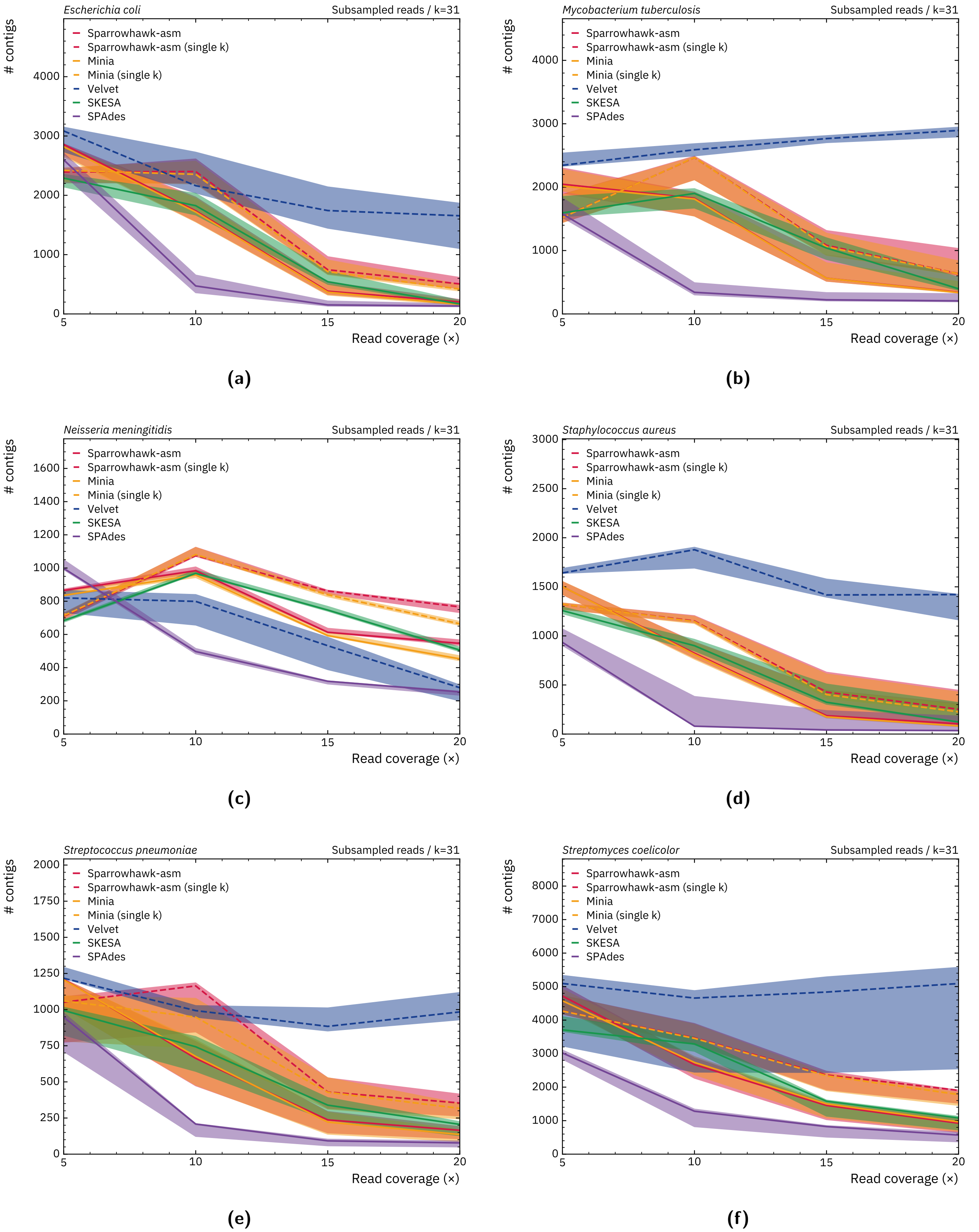


**Supplementary fig. 9.** `sparrowhawk-asm` benchmark on subsampled real reads: number of contigs per species against coverage (three read sets each). Lines show the median over read sets at each subsampled coverage, and shaded bands the interquartile range. Single-$k$ assemblers ran at $k = 31$. **a,** *Escherichia coli*; **b,** *Mycobacterium tuberculosis*; **c,** *Neisseria meningitidis*; **d,** *Staphylococcus aureus*; **e,** *Streptococcus pneumoniae*; **f,** *Streptomyces coelicolor*.

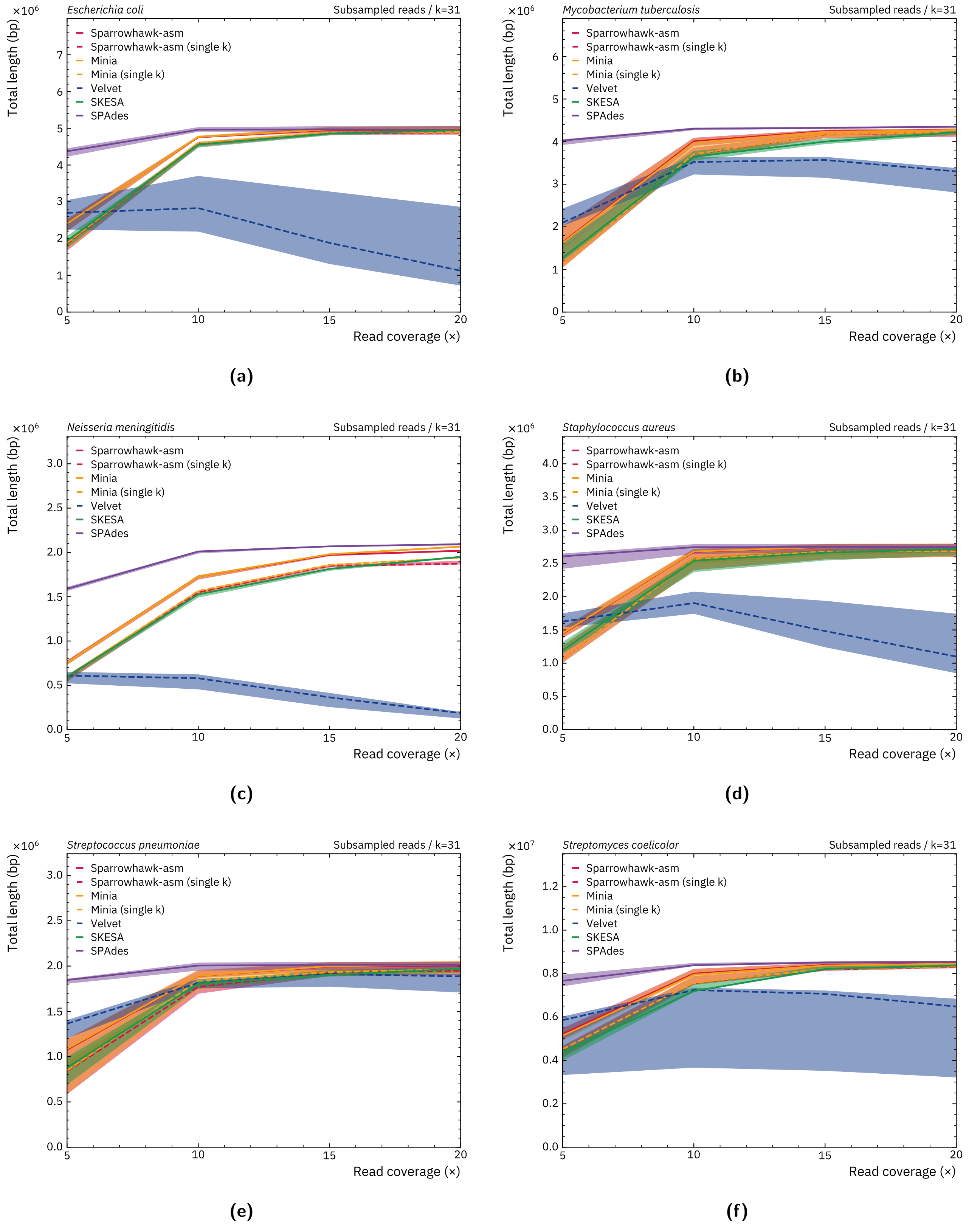


**Supplementary fig. 10.** `sparrowhawk-asm` benchmark on subsampled real reads: total assembly length per species against coverage (three read sets each). Lines show the median over read sets at each subsampled coverage, and shaded bands the interquartile range. Single-$k$ assemblers ran at $k = 31$. **a,** *Escherichia coli*; **b,** *Mycobacterium tuberculosis*; **c,** *Neisseria meningitidis*; **d,** *Staphylococcus aureus*; **e,** *Streptococcus pneumoniae*; **f,** *Streptomyces coelicolor*.

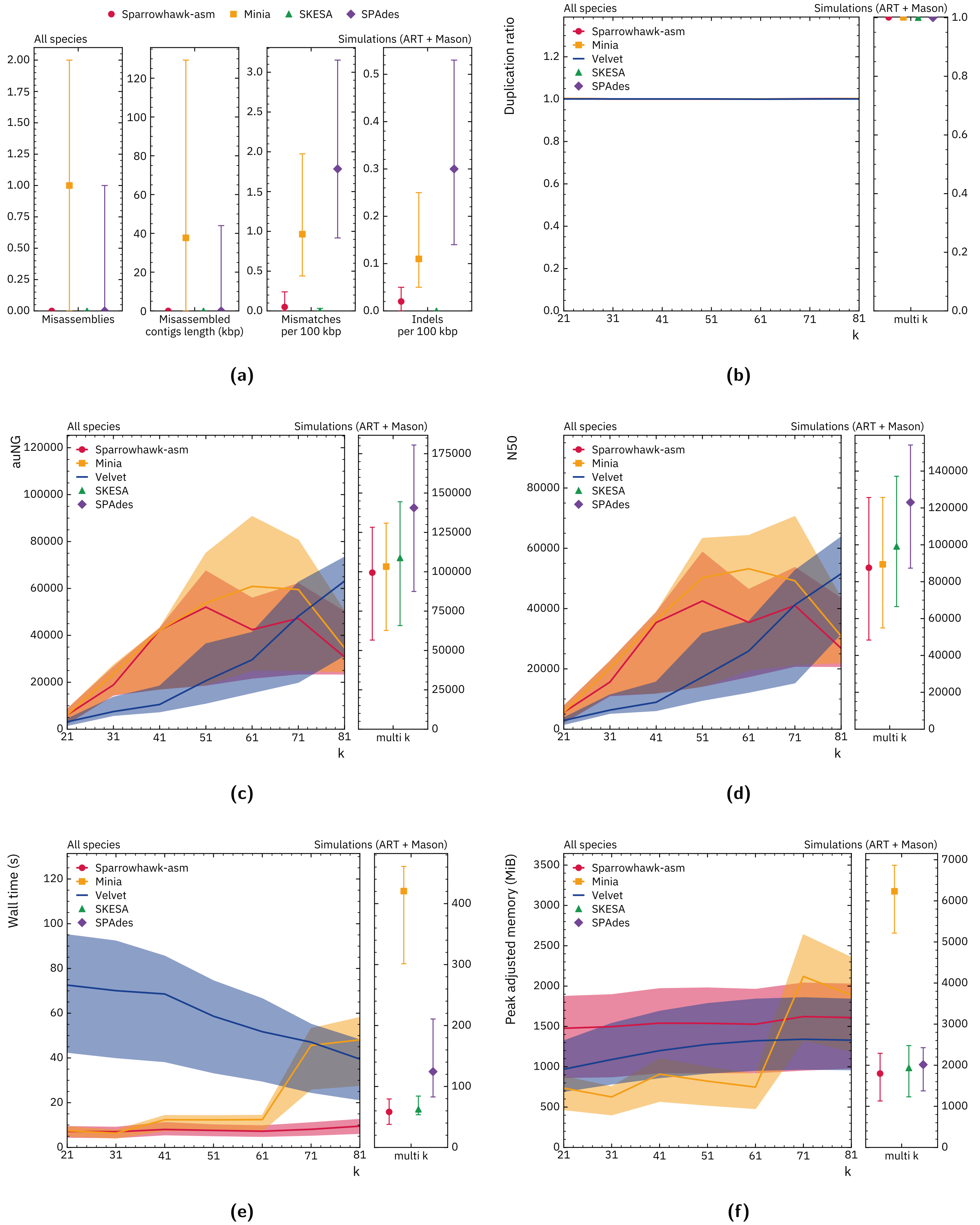


**Supplementary fig. 11.** `sparrowhawk-asm` benchmark on reads simulated with `ART` and `Mason`, pooled over the six species: **a,** Number of misassemblies, misassembled contigs length, and mismatches and indels per 100 kbp; **b,** Duplication ratio; **c,** auNG; **d,** N50; **e,** Wall time; **f,** Peak adjusted memory. **a** shows only the multi-$k$ assemblers: Points show the median over read sets, and whiskers the interquartile range. In **b**–**f**: Lines show the median over read sets at each $k$-mer length, and shaded bands the interquartile range. The right-hand panel shows the multi-$k$ assemblers (`sparrowhawk-asm`, `Minia`, `SKESA`, and `SPAdes`) as medians with interquartile whiskers, on their own $y$ axis. Peak adjusted memory includes both RAM as well as the file disk allocation. We use for this the peak memory charged to the run's control group (cgroup), which includes the in-memory (tmpfs) working directory used by `Minia` and `SPAdes`.

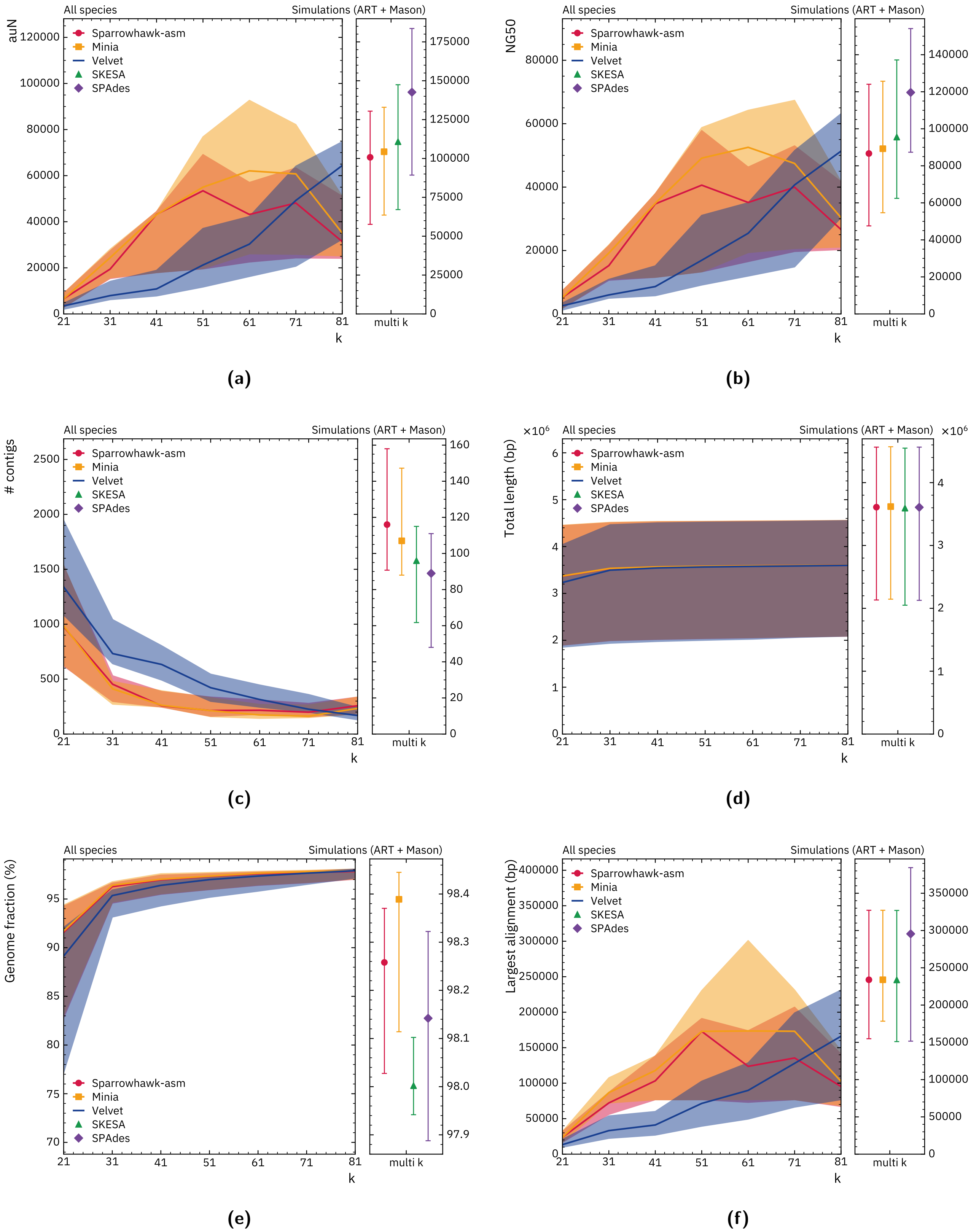


**Supplementary fig. 12.** `sparrowhawk-asm` benchmark on reads simulated with `ART` and `Mason`, pooled over the six species: **a,** auN; **b,** NG50; **c,** Number of contigs; **d,** Total length; **e,** Genome fraction; **f,** Largest alignment. Lines show the median over read sets at each $k$-mer length, and shaded bands the interquartile range. The right-hand panel shows the multi-$k$ assemblers (`sparrowhawk-asm`, `Minia`, `SKESA`, and `SPAdes`) as medians with interquartile whiskers, on their own $y$ axis.

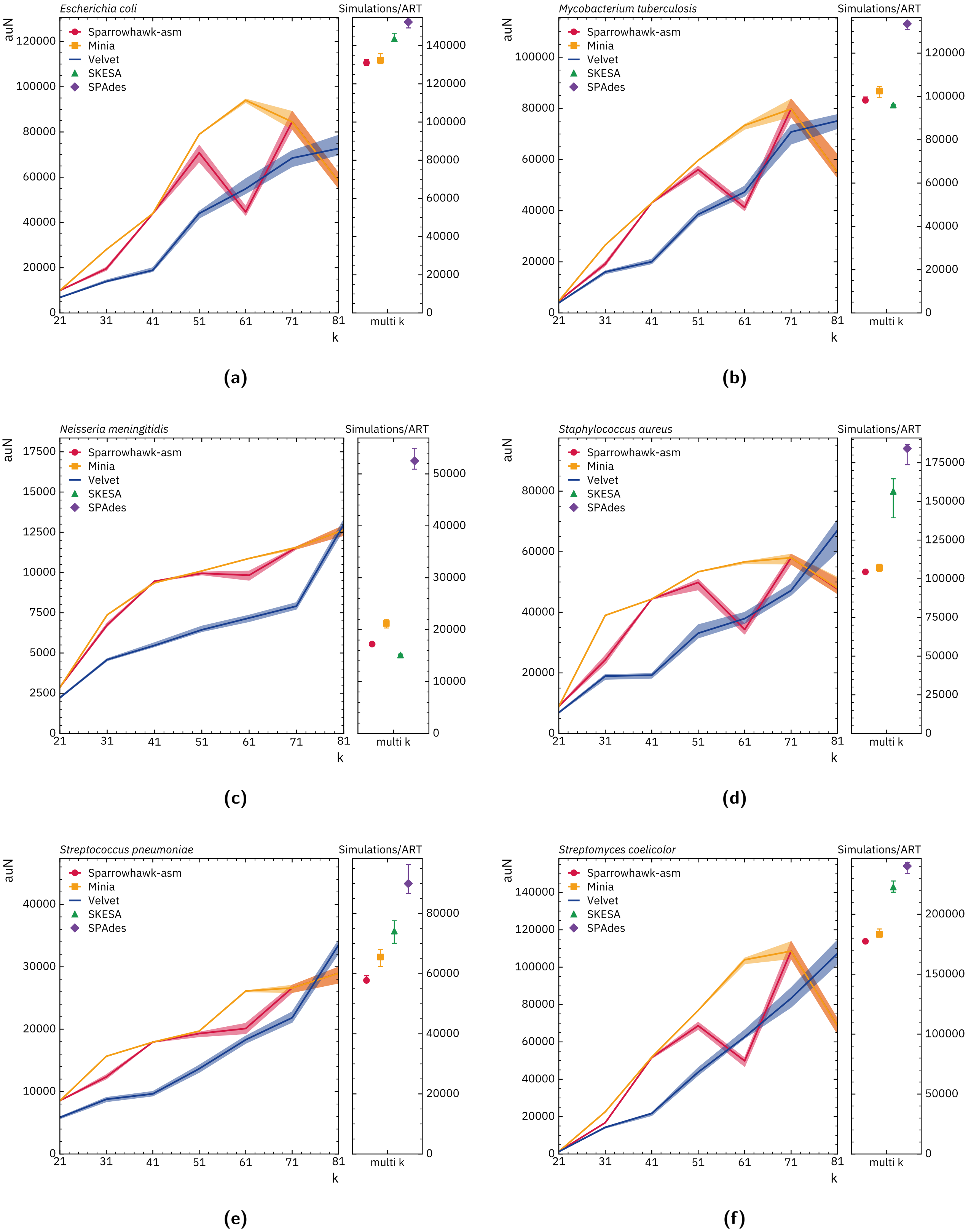


**Supplementary fig. 13.** `sparrowhawk-asm` benchmark on reads simulated with `ART`: auN per species. Lines show the median over read sets at each $k$-mer length, and shaded bands the interquartile range. The right-hand panel shows the multi-$k$ assemblers (`sparrowhawk-asm`, `Minia`, `SKESA`, and `SPAdes`) as medians with interquartile whiskers, on their own $y$ axis. **a,** *Escherichia coli*; **b,** *Mycobacterium tuberculosis*; **c,** *Neisseria meningitidis*; **d,** *Staphylococcus aureus*; **e,** *Streptococcus pneumoniae*; **f,** *Streptomyces coelicolor*.

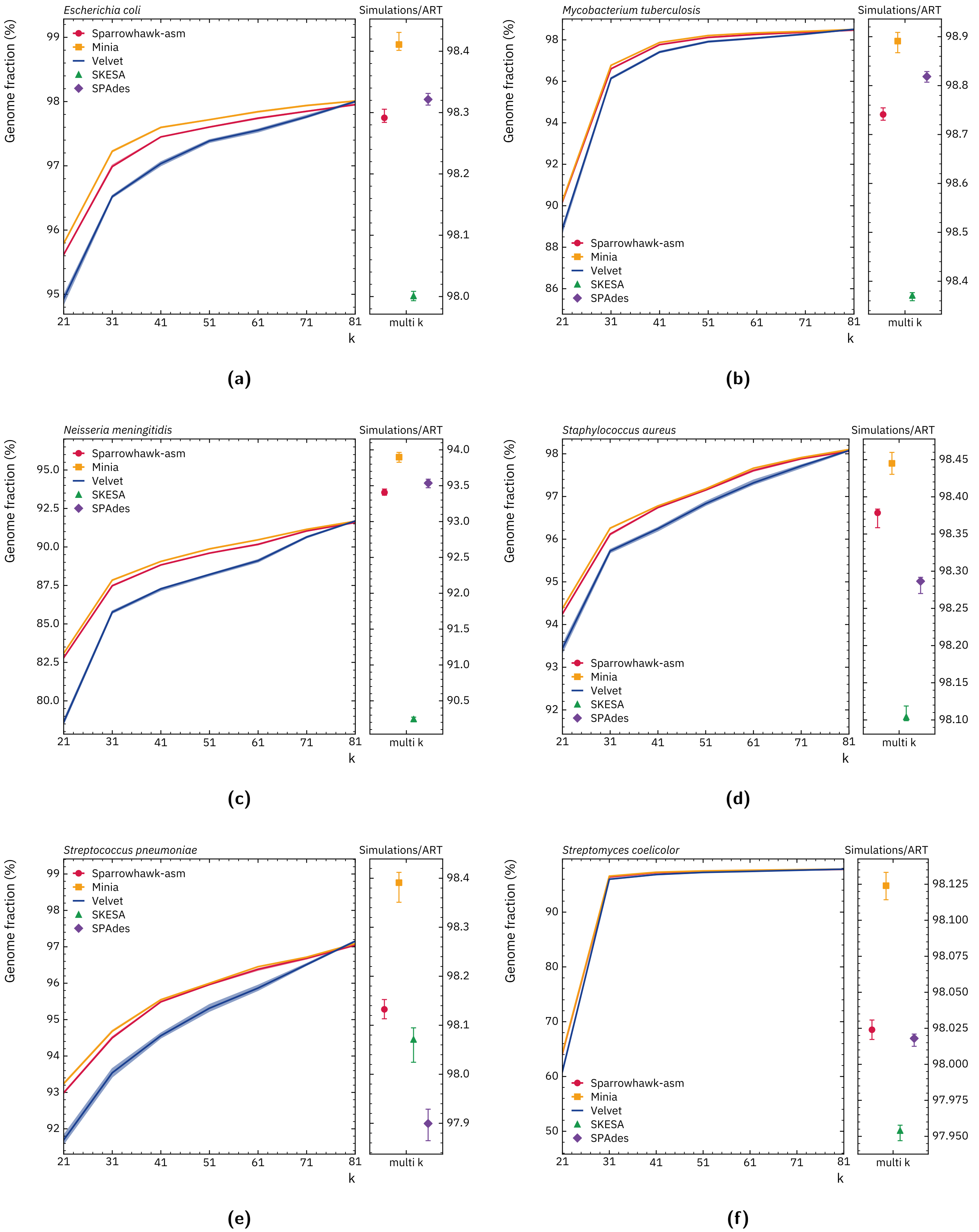


**Supplementary fig. 14.** `sparrowhawk-asm` benchmark on reads simulated with `ART`: genome fraction per species. Lines show the median over read sets at each $k$-mer length, and shaded bands the interquartile range. The right-hand panel shows the multi-$k$ assemblers (`sparrowhawk-asm`, `Minia`, `SKESA`, and `SPAdes`) as medians with interquartile whiskers, on their own $y$ axis. **a,** *Escherichia coli*; **b,** *Mycobacterium tuberculosis*; **c,** *Neisseria meningitidis*; **d,** *Staphylococcus aureus*; **e,** *Streptococcus pneumoniae*; **f,** *Streptomyces coelicolor*.

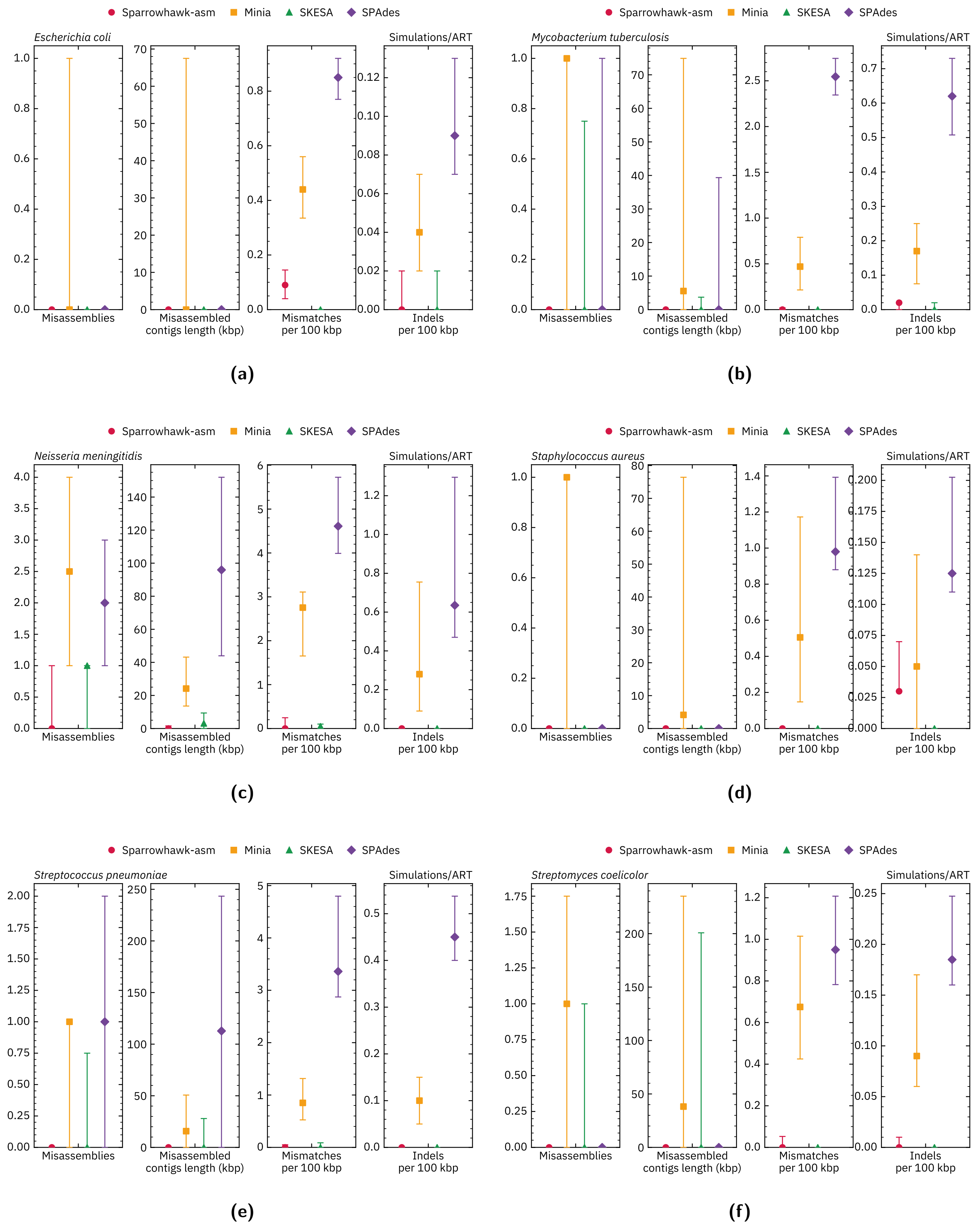


**Supplementary fig. 15.** `sparrowhawk-asm` benchmark on reads simulated with `ART`: number of misassemblies, misassembled contigs length, and mismatches and indels per 100 kbp, per species, for the multi-$k$ assemblers. Points show the median over read sets, and whiskers the interquartile range. **a,** *Escherichia coli*; **b,** *Mycobacterium tuberculosis*; **c,** *Neisseria meningitidis*; **d,** *Staphylococcus aureus*; **e,** *Streptococcus pneumoniae*; **f,** *Streptomyces coelicolor*.

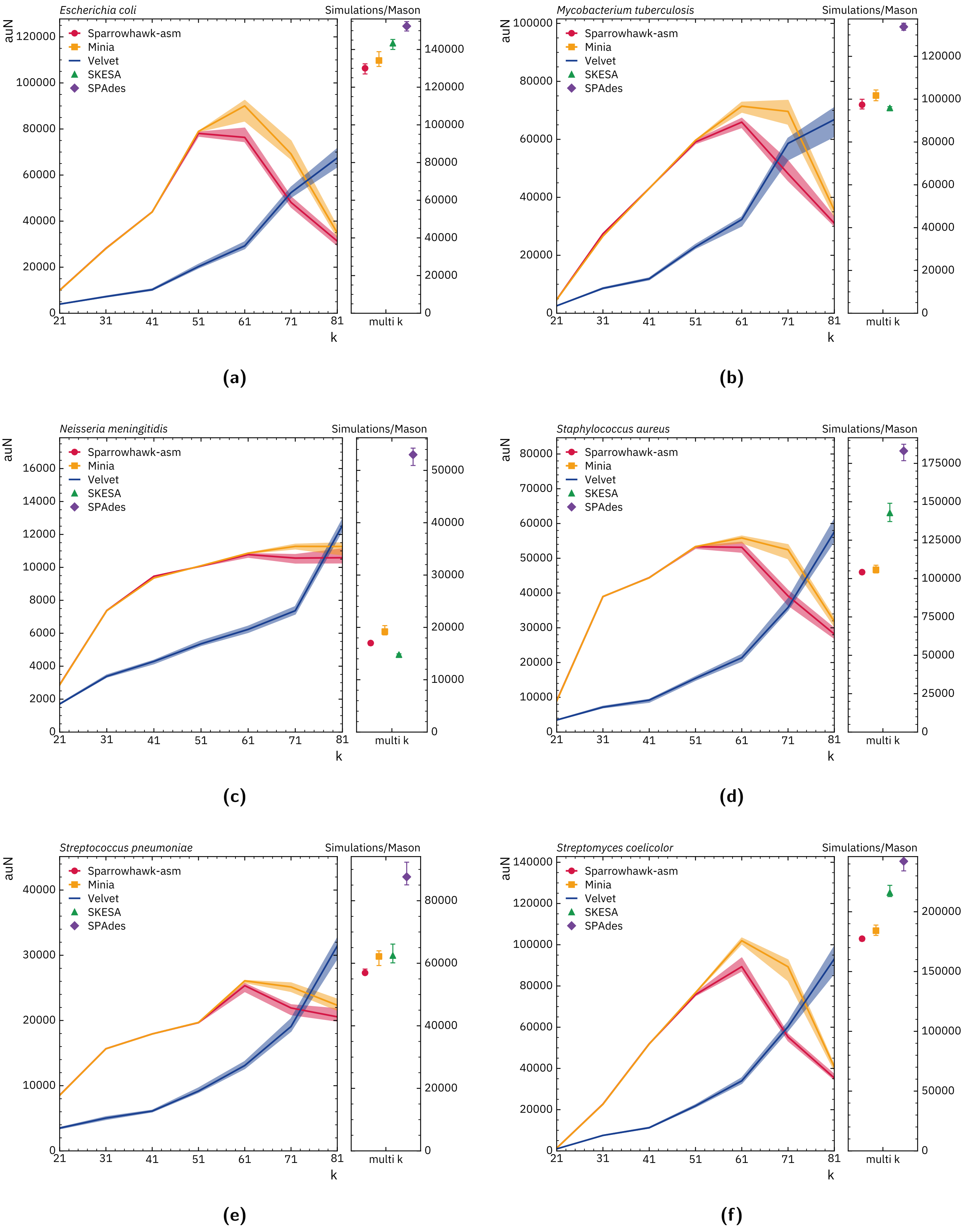


**Supplementary fig. 16.** `sparrowhawk-asm` benchmark on reads simulated with `Mason`: auN per species. Lines show the median over read sets at each $k$-mer length, and shaded bands the interquartile range. The right-hand panel shows the multi-$k$ assemblers (`sparrowhawk-asm`, `Minia`, `SKESA`, and `SPAdes`) as medians with interquartile whiskers, on their own $y$ axis. **a,** *Escherichia coli*; **b,** *Mycobacterium tuberculosis*; **c,** *Neisseria meningitidis*; **d,** *Staphylococcus aureus*; **e,** *Streptococcus pneumoniae*; **f,** *Streptomyces coelicolor*.

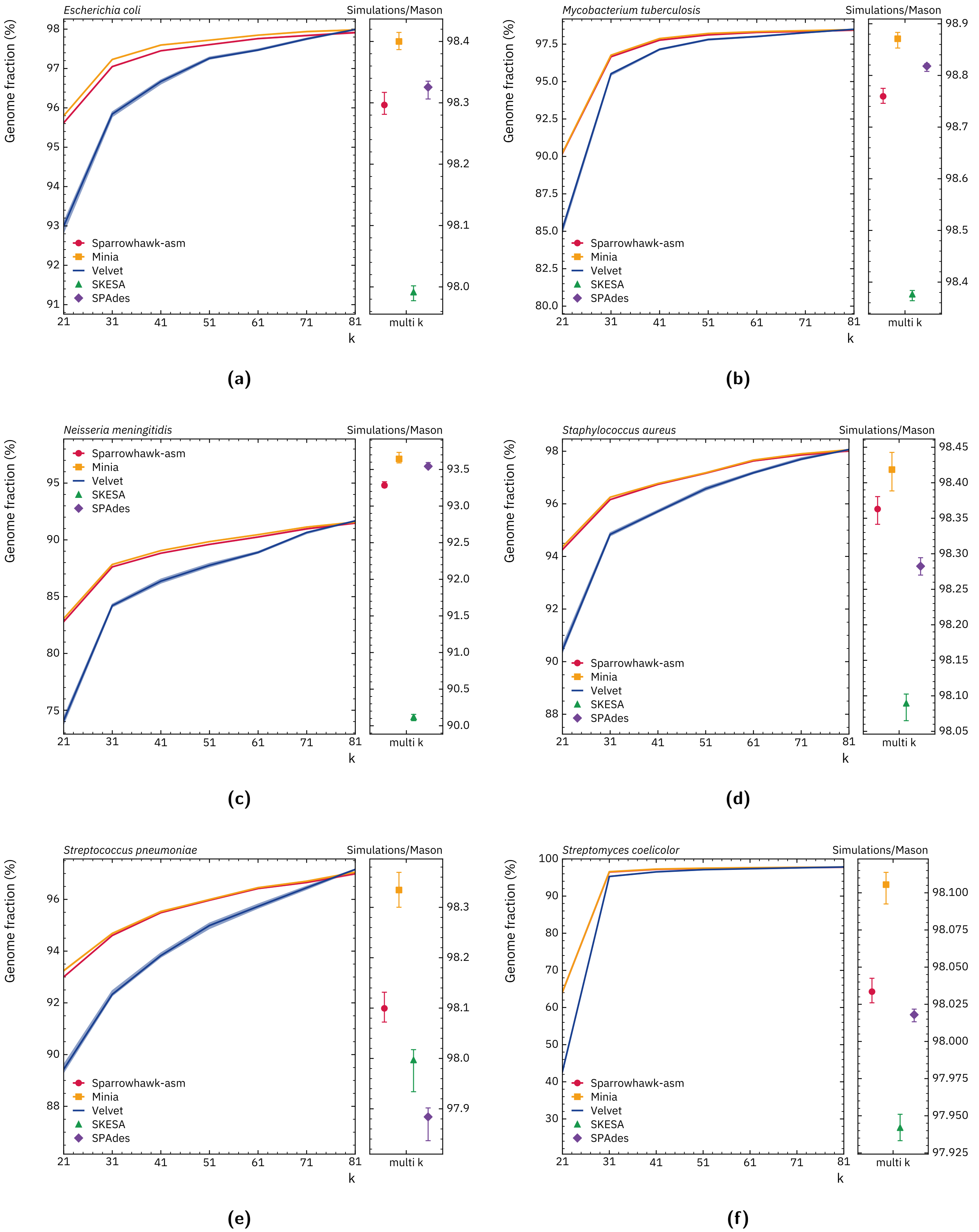


**Supplementary fig. 17.** `sparrowhawk-asm` benchmark on reads simulated with `Mason`: genome fraction per species. Lines show the median over read sets at each $k$-mer length, and shaded bands the interquartile range. The right-hand panel shows the multi-$k$ assemblers (`sparrowhawk-asm`, `Minia`, `SKESA`, and `SPAdes`) as medians with interquartile whiskers, on their own $y$ axis. **a,** *Escherichia coli*; **b,** *Mycobacterium tuberculosis*; **c,** *Neisseria meningitidis*; **d,** *Staphylococcus aureus*; **e,** *Streptococcus pneumoniae*; **f,** *Streptomyces coelicolor*.

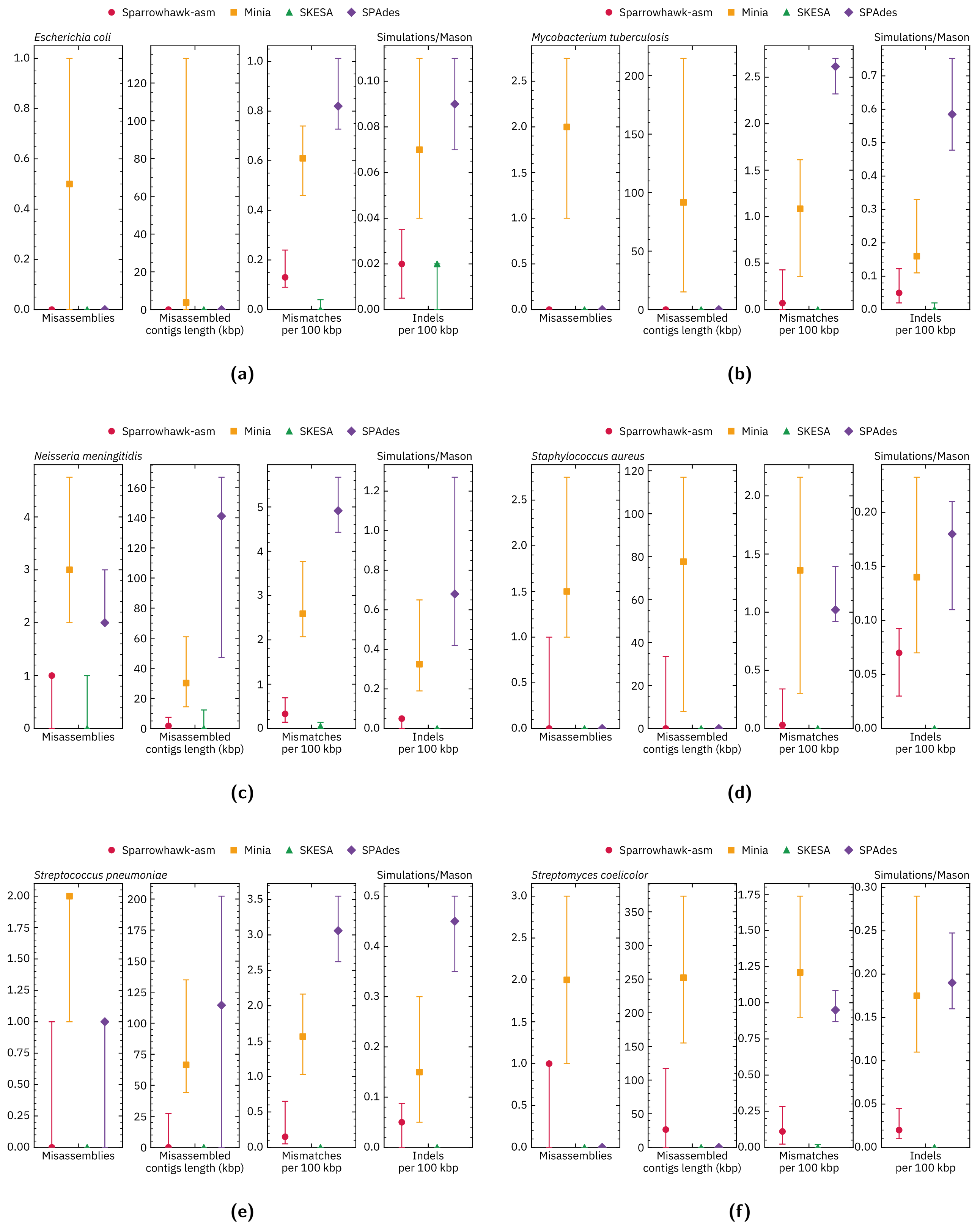


**Supplementary fig. 18.** `sparrowhawk-asm` benchmark on reads simulated with `Mason`: number of misassemblies, misassembled contigs length, and mismatches and indels per 100 kbp, per species, for the multi-$k$ assemblers. Points show the median over read sets, and whiskers the interquartile range. **a,** *Escherichia coli*; **b,** *Mycobacterium tuberculosis*; **c,** *Neisseria meningitidis*; **d,** *Staphylococcus aureus*; **e,** *Streptococcus pneumoniae*; **f,** *Streptomyces coelicolor*.

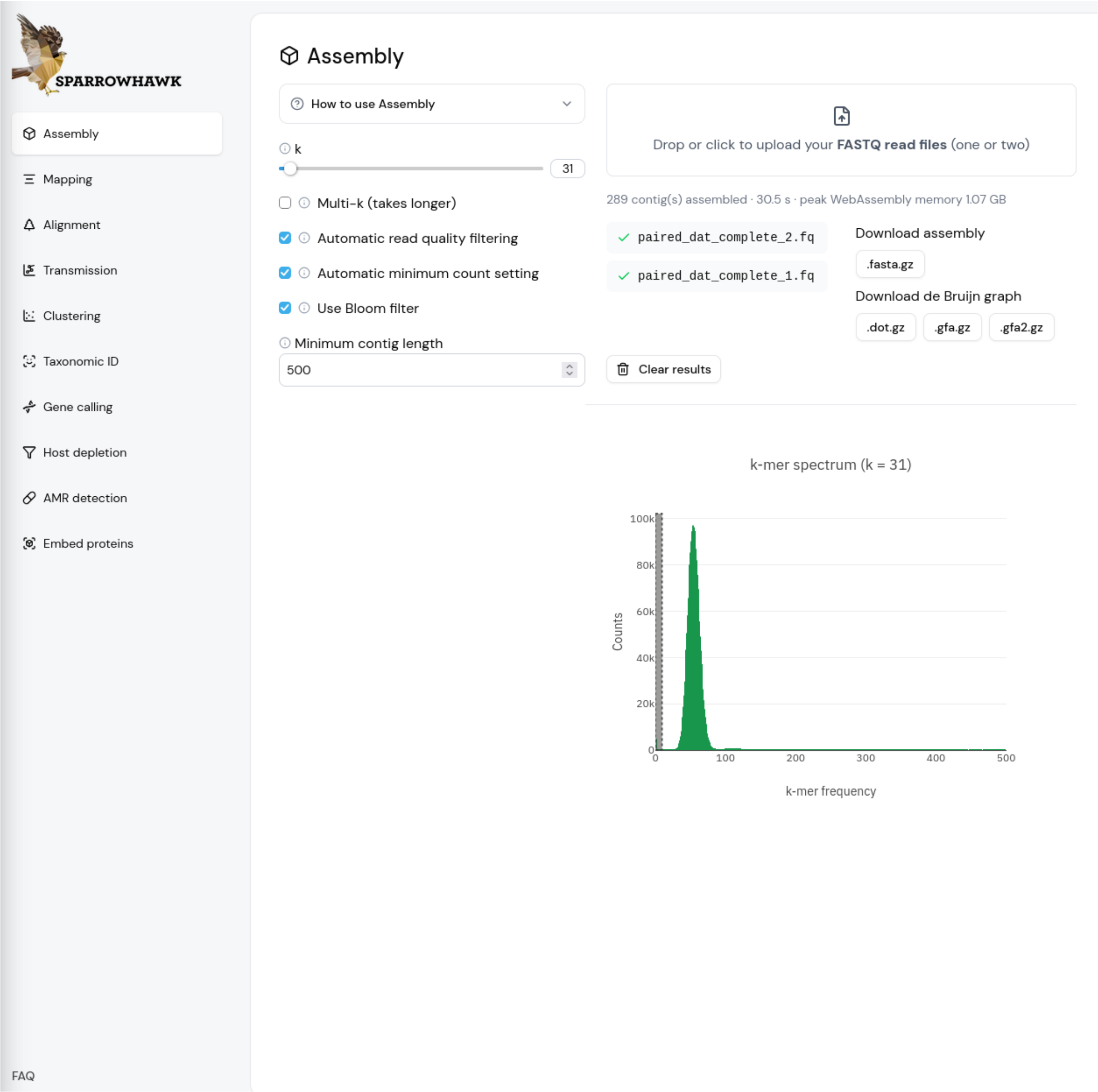


**Supplementary fig. 19.** Screenshot of `sparrowhawk` on the "Assembly" tab.

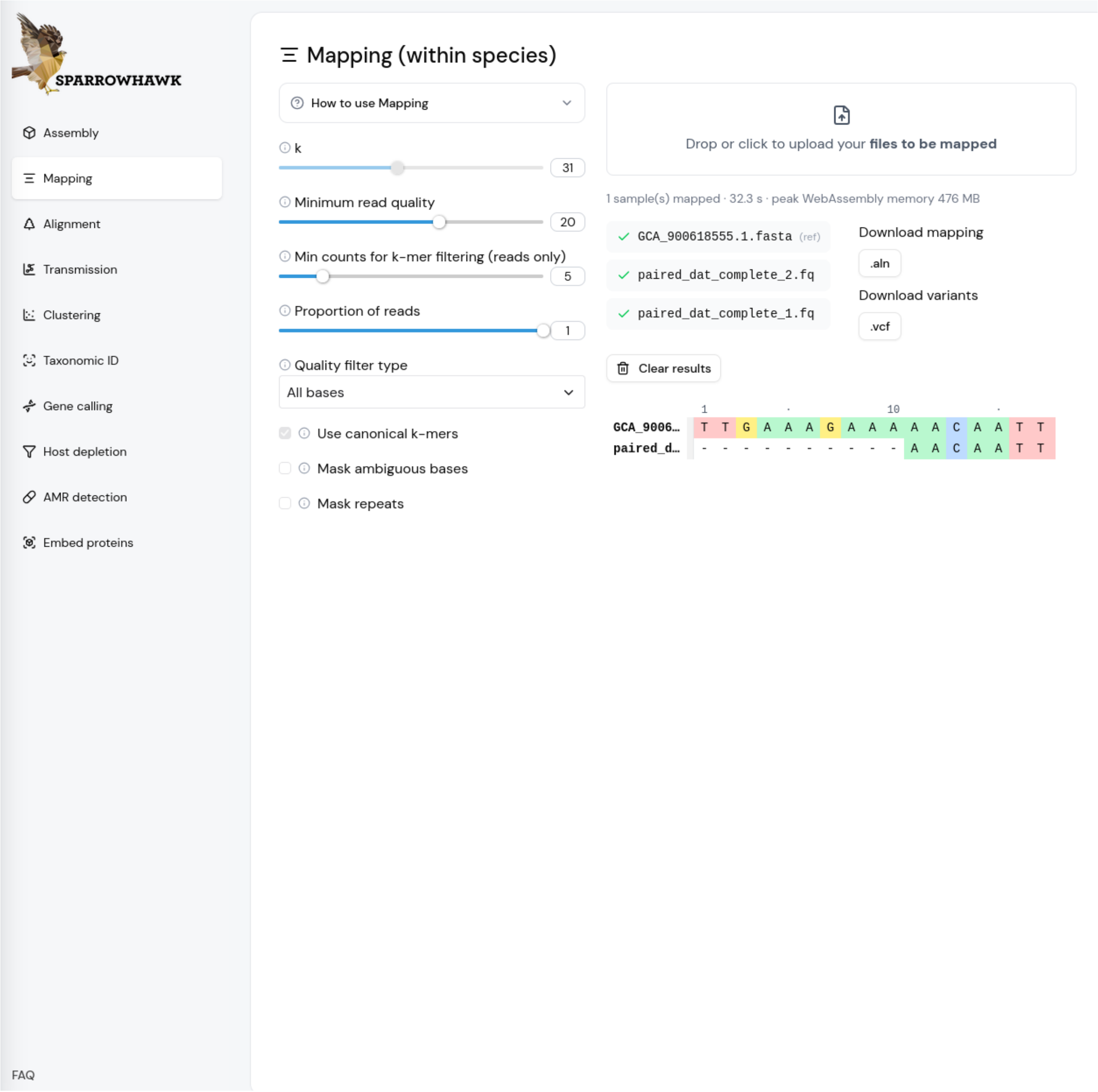


**Supplementary fig. 20.** Screenshot of `sparrowhawk` on the "Mapping" tab.

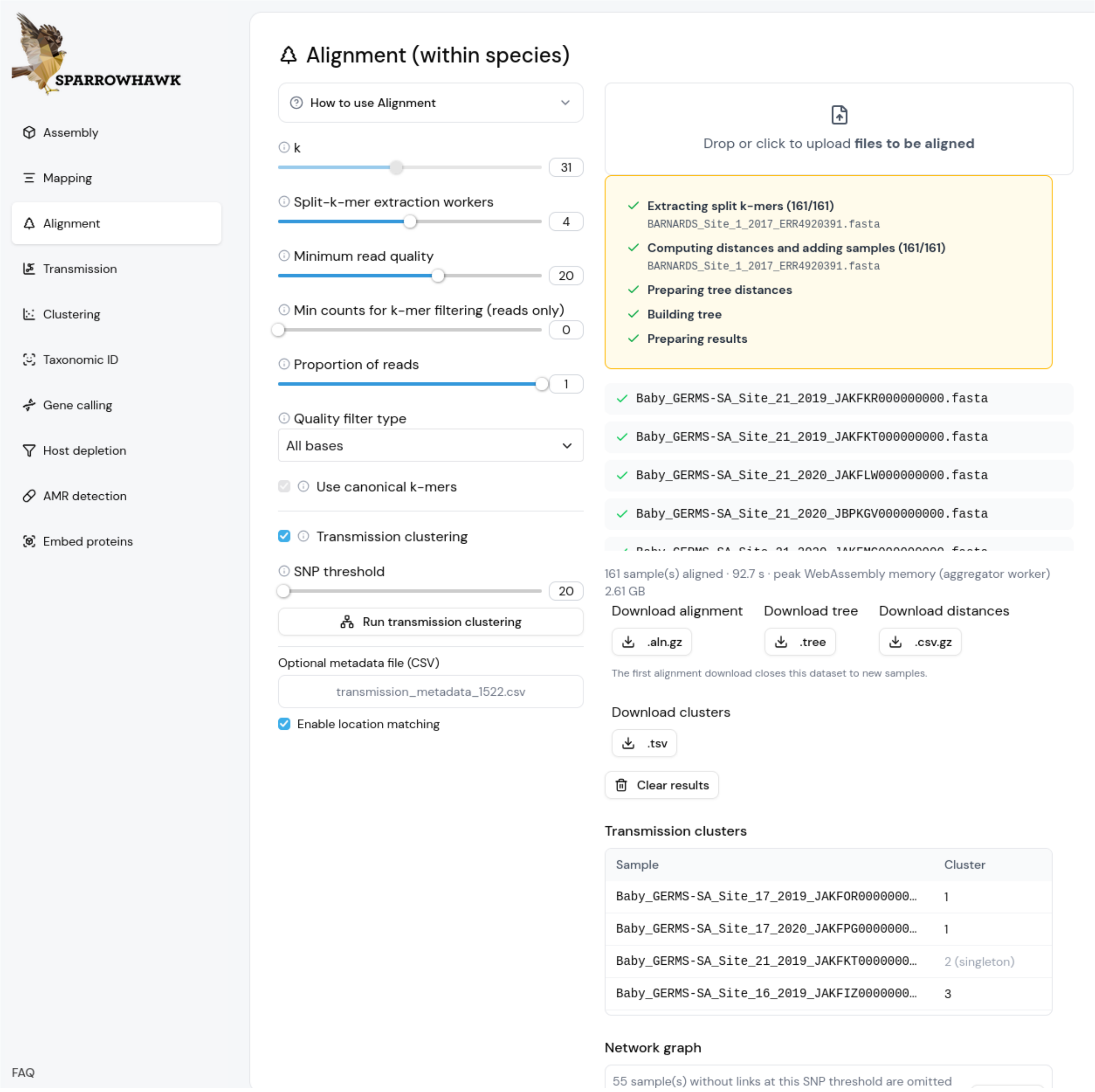


**Supplementary fig. 21.** Screenshot of `sparrowhawk` on the "Alignment" tab, top part.

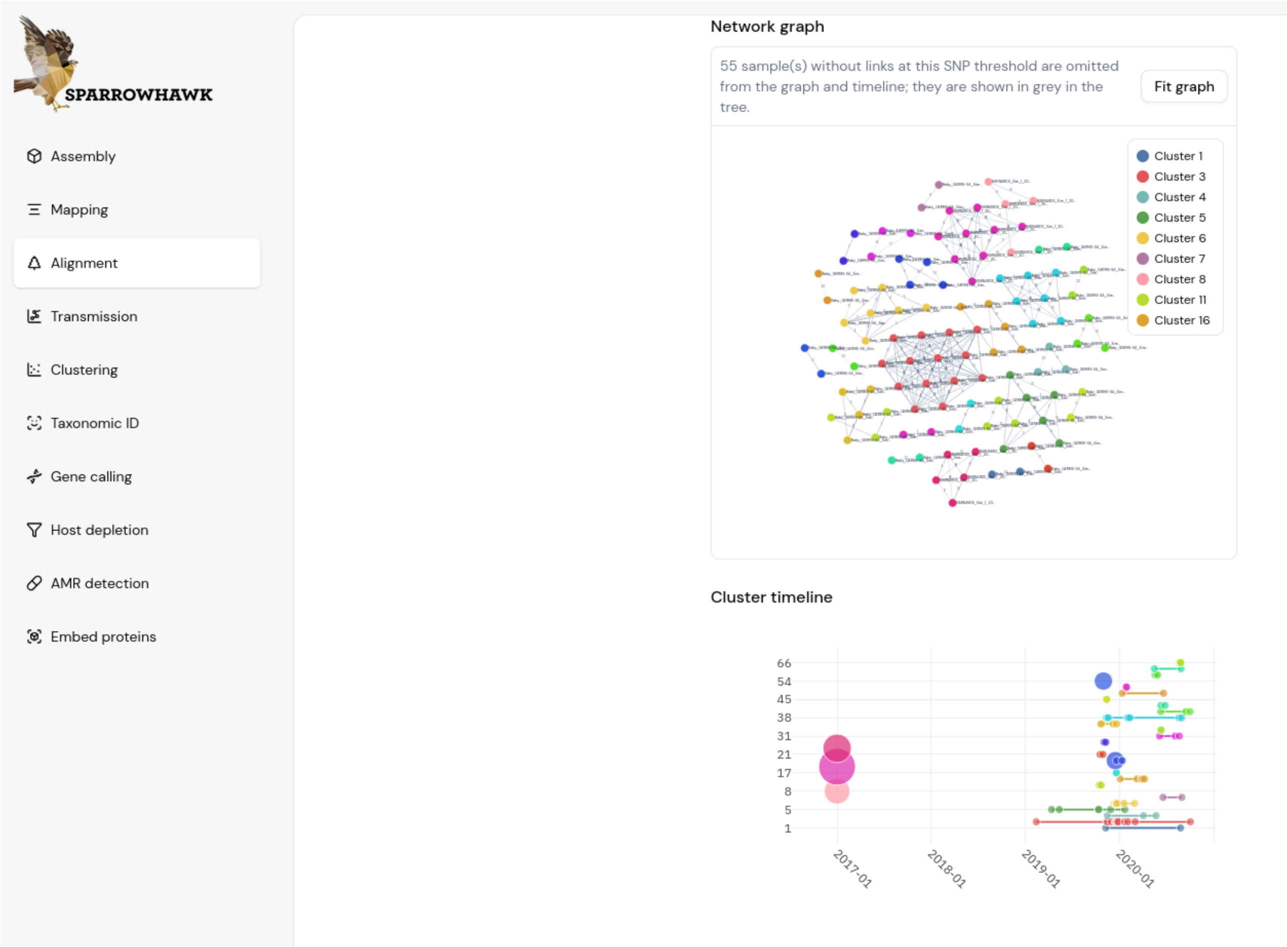


**Supplementary fig. 22.** Screenshot of `sparrowhawk` on the "Alignment" tab, middle part.

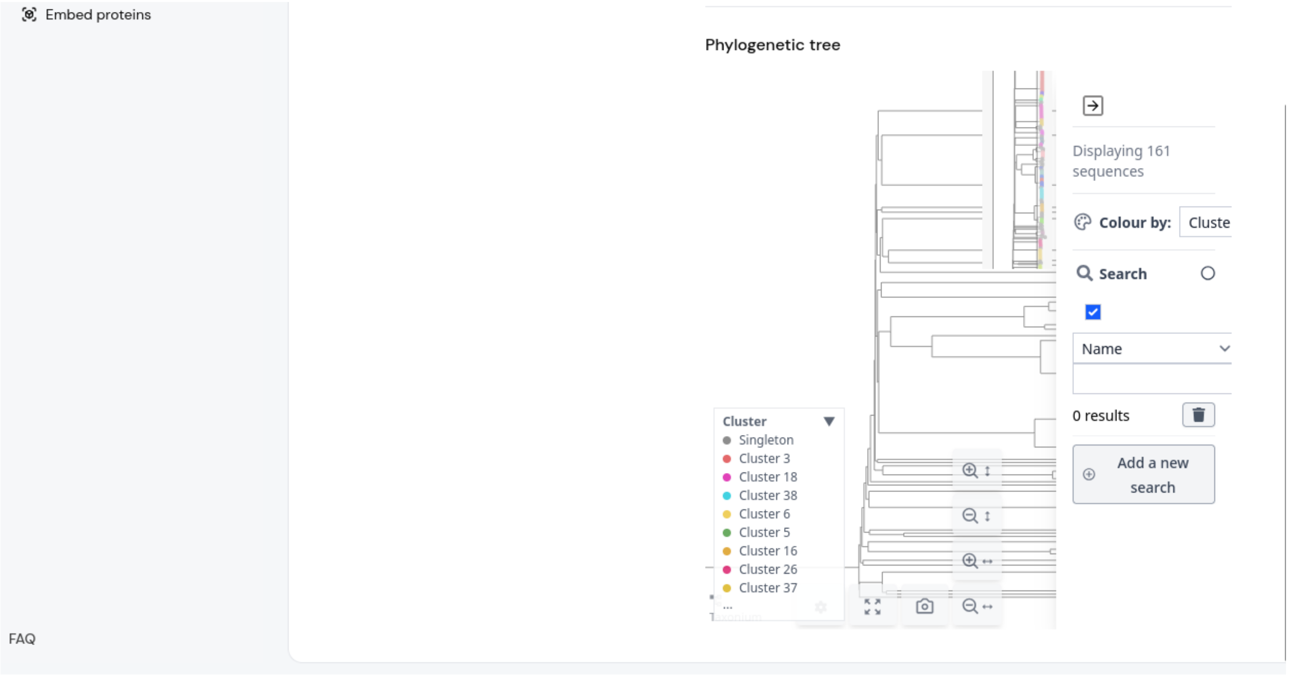


**Supplementary fig. 23.** Screenshot of `sparrowhawk` on the "Alignment" tab, bottom part.

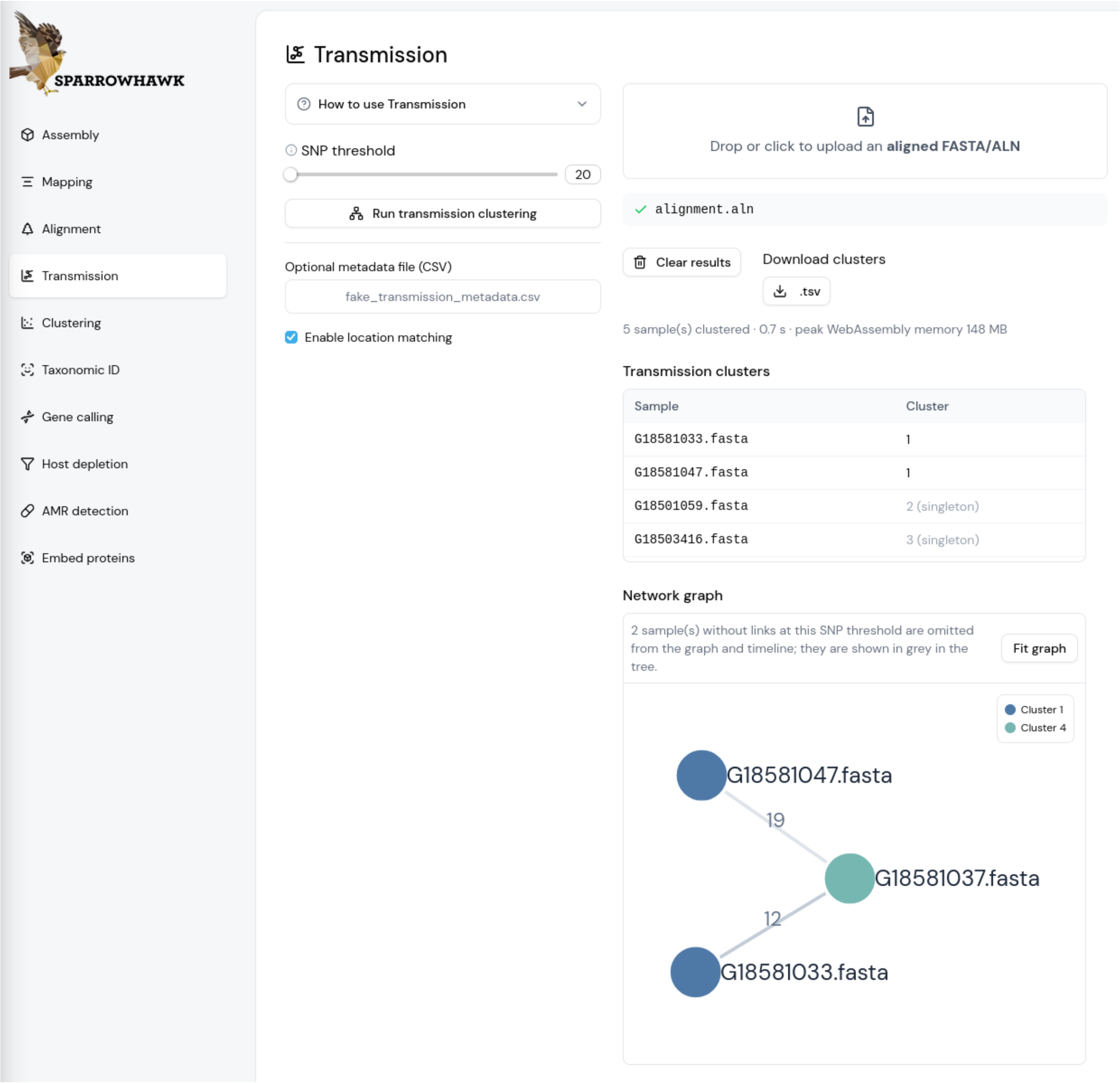


**Supplementary fig. 24.** Screenshot of `sparrowhawk` on the "Transmission" tab.

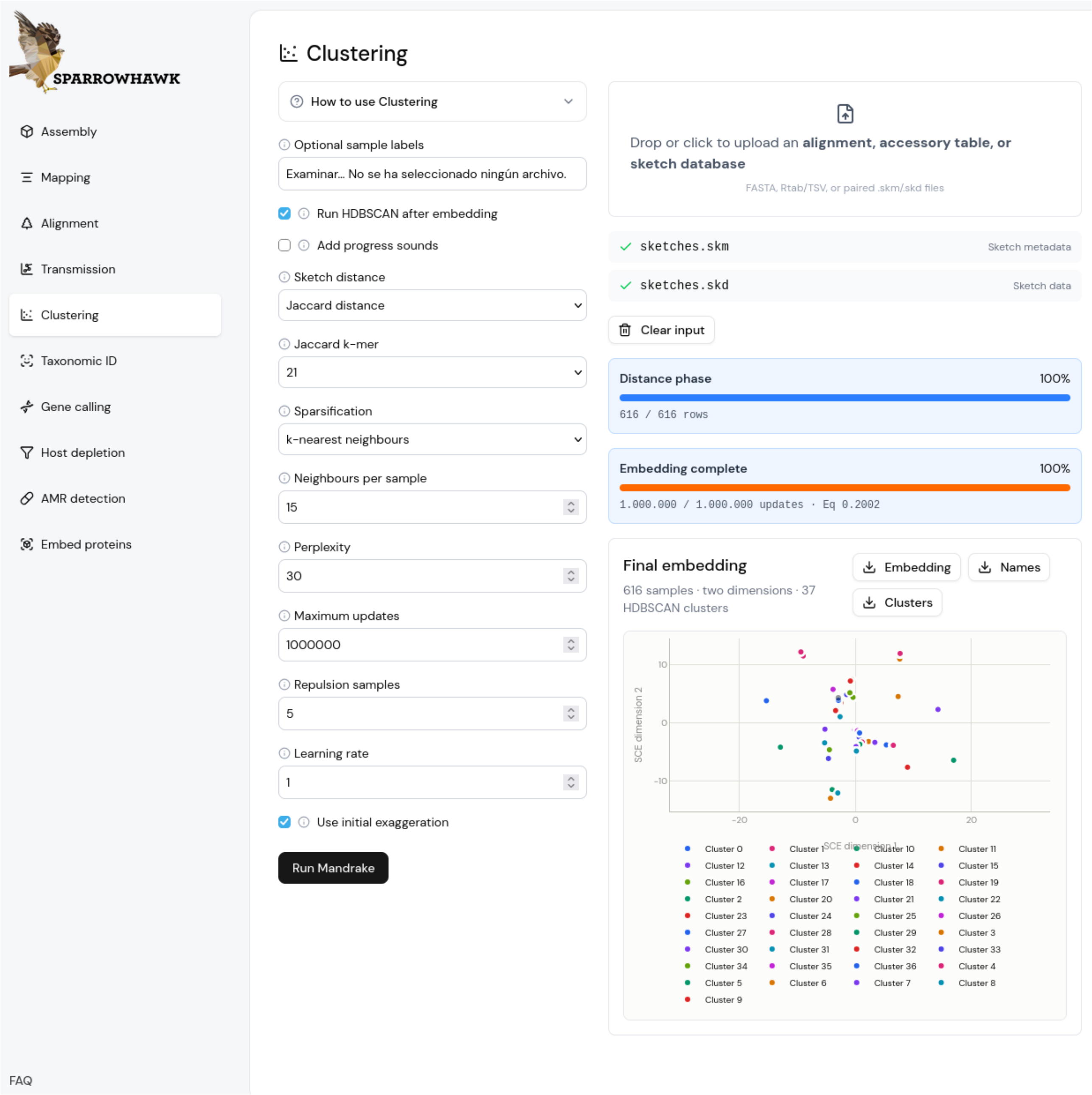


**Supplementary fig. 25.** Screenshot of `sparrowhawk` on the “Clustering” tab.

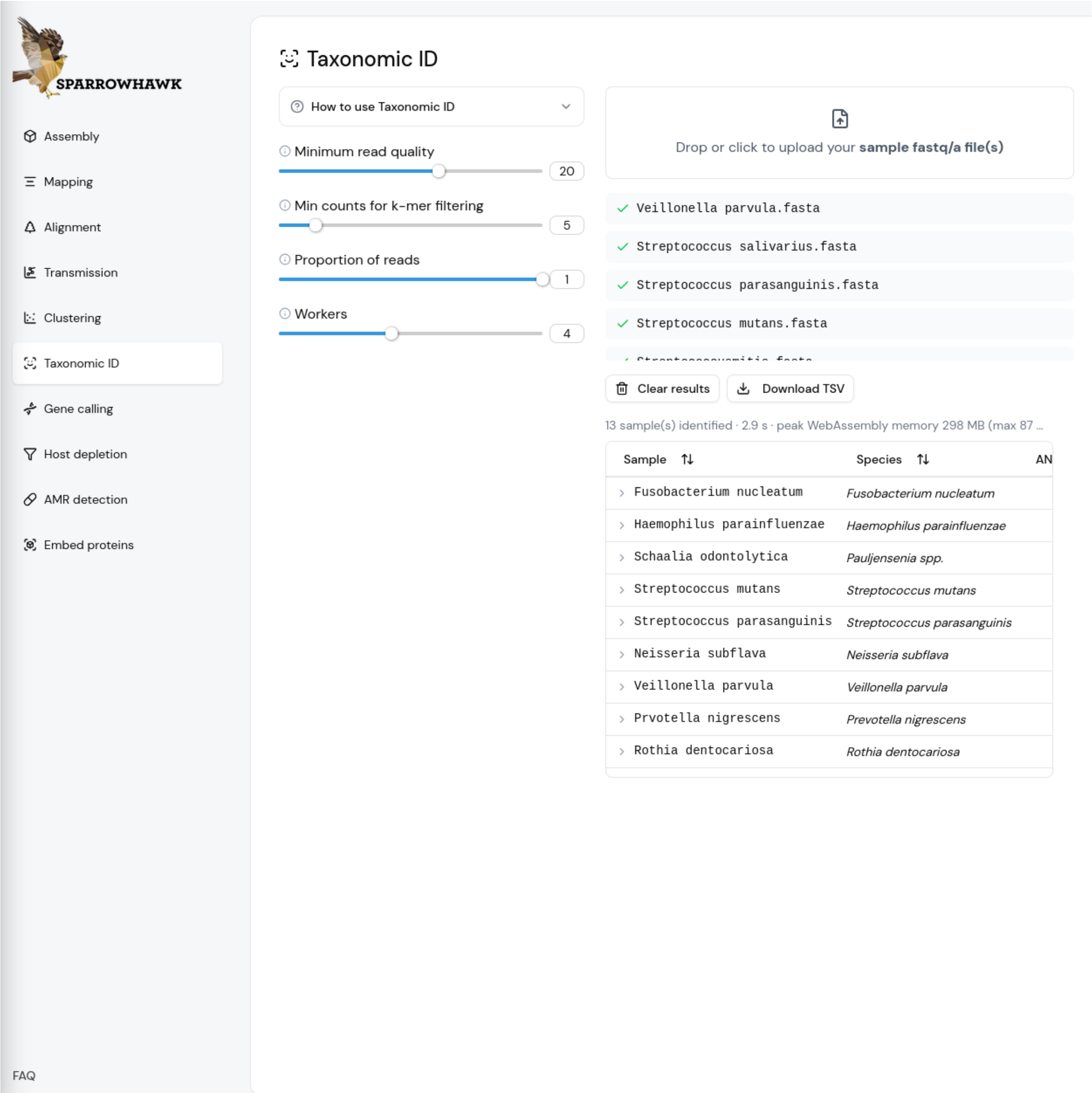

**Supplementary fig. 26.** Screenshot of `sparrowhawk` on the "Taxonomic ID" tab.

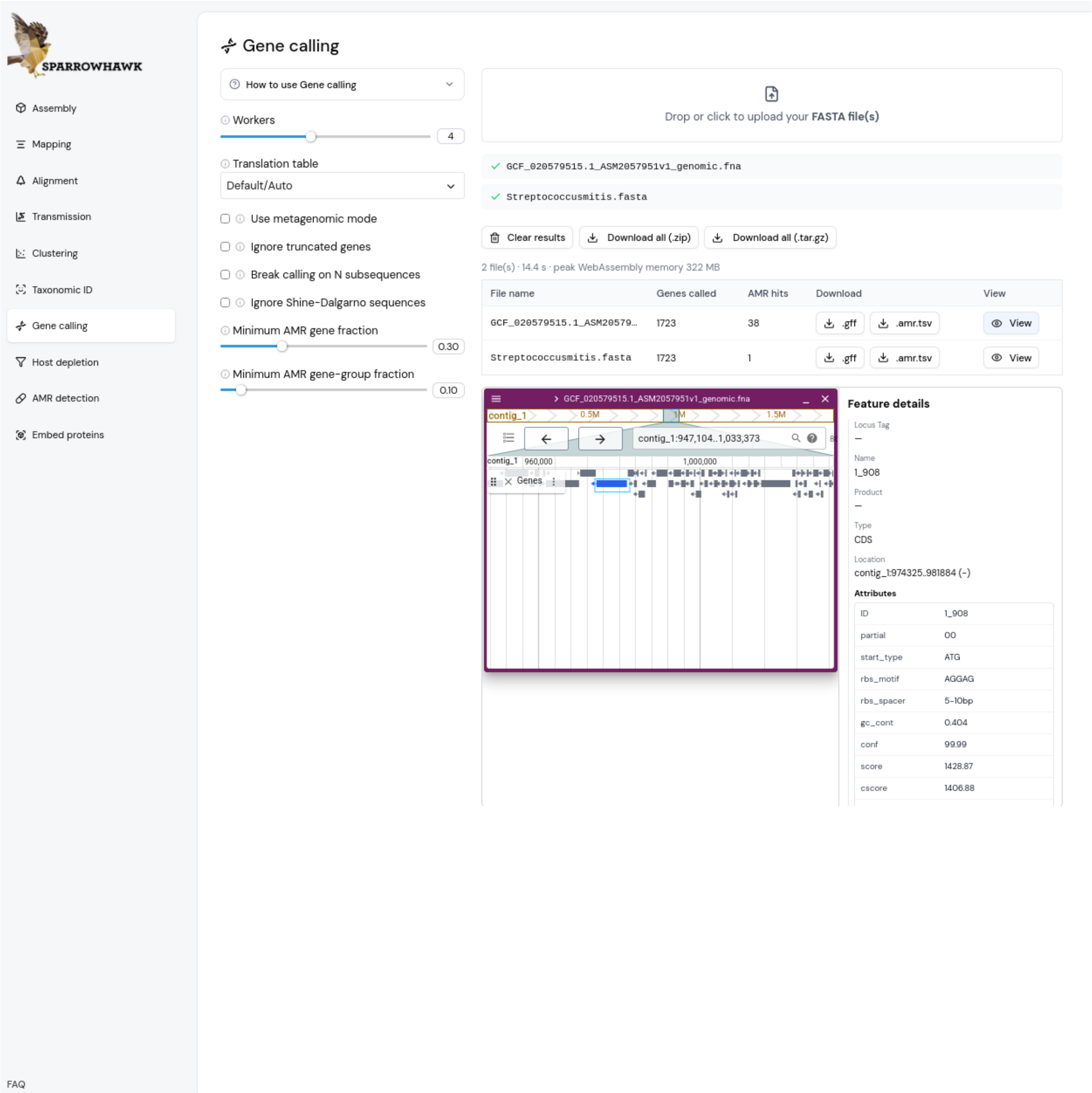


**Supplementary fig. 27.** Screenshot of `sparrowhawk` on the "Gene calling" tab.

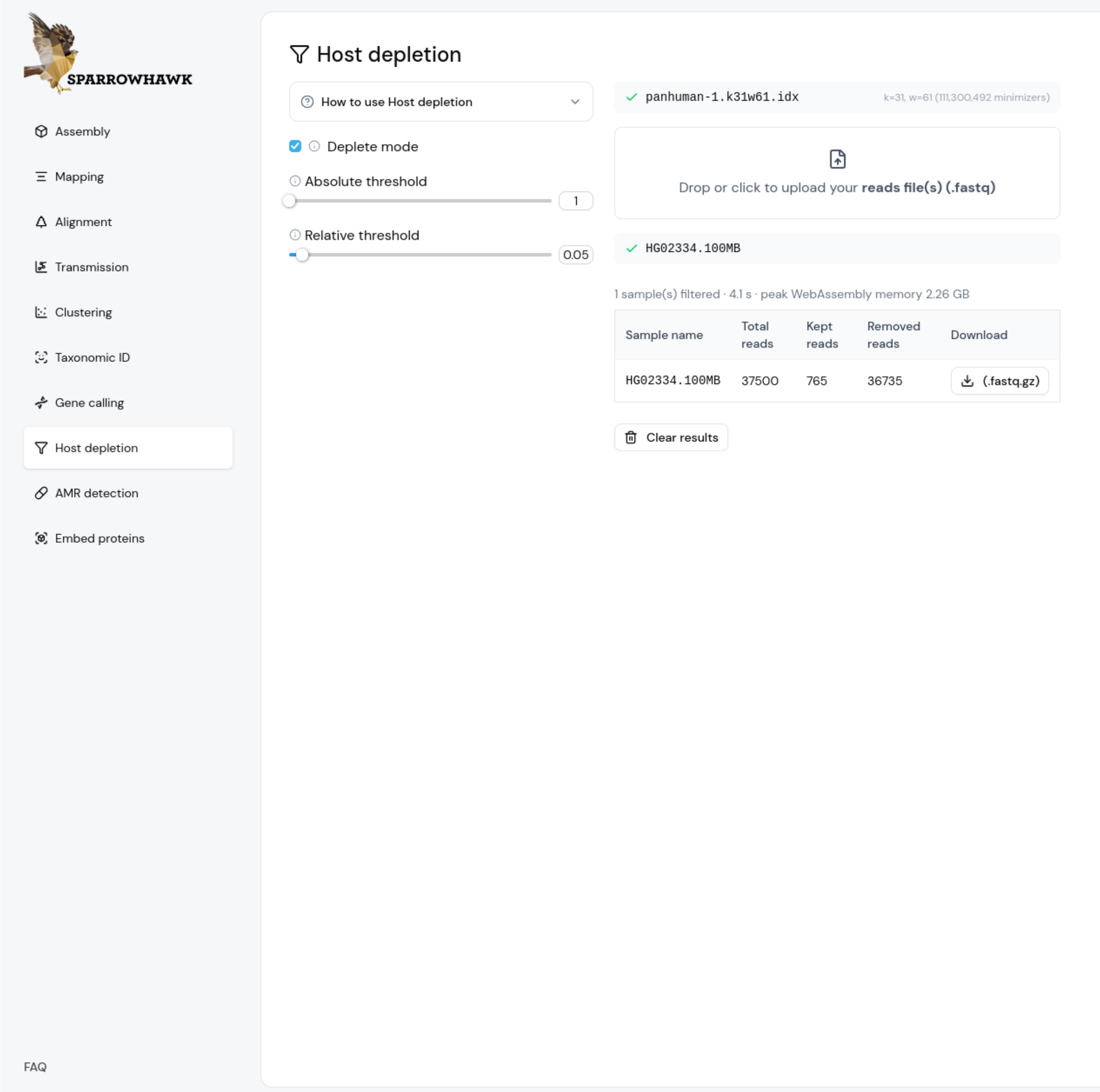


**Supplementary fig. 28.** Screenshot of `sparrowhawk` on the "Host depletion" tab.

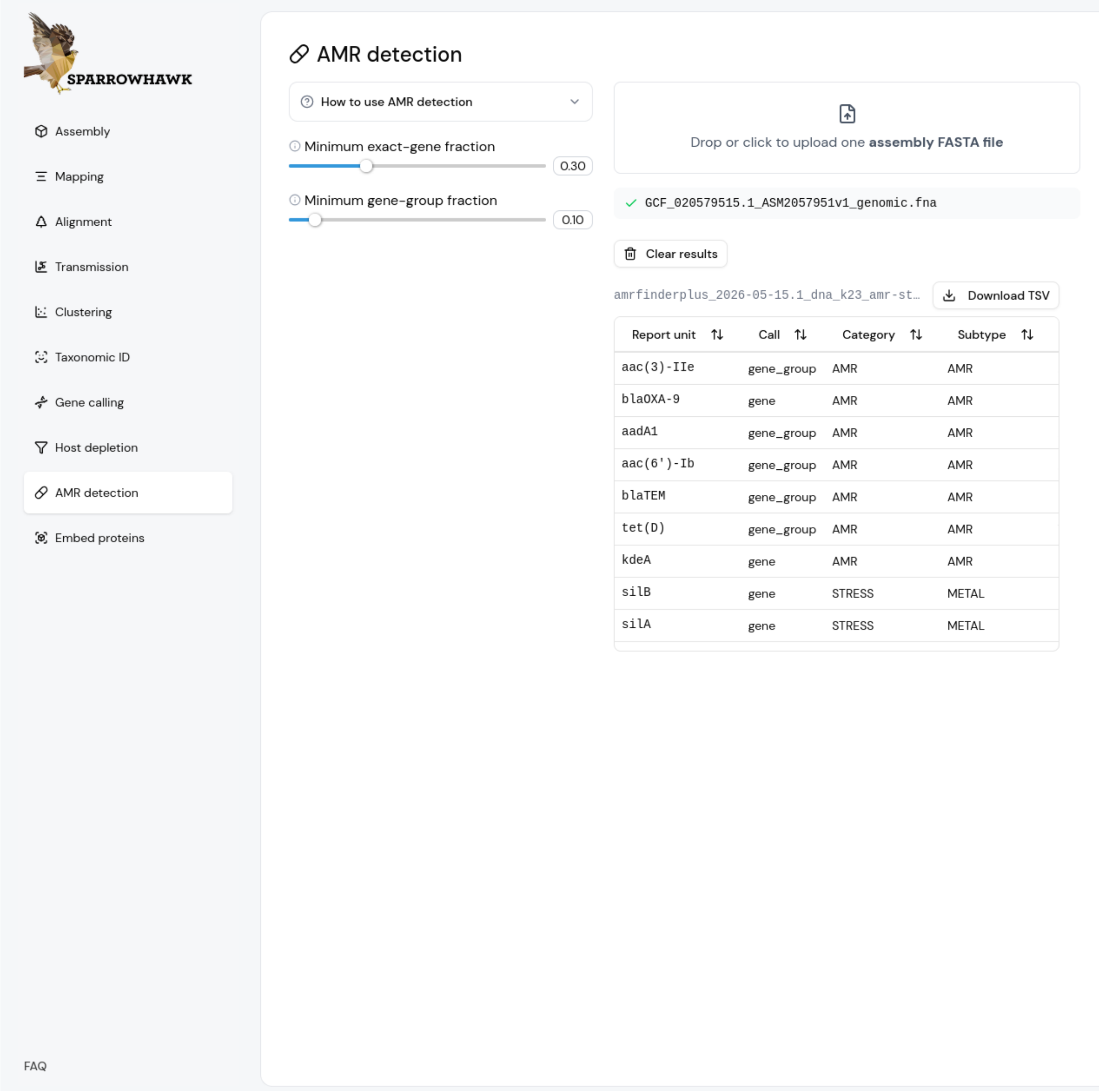


**Supplementary fig. 29.** Screenshot of `sparrowhawk` on the "AMR detection" tab.

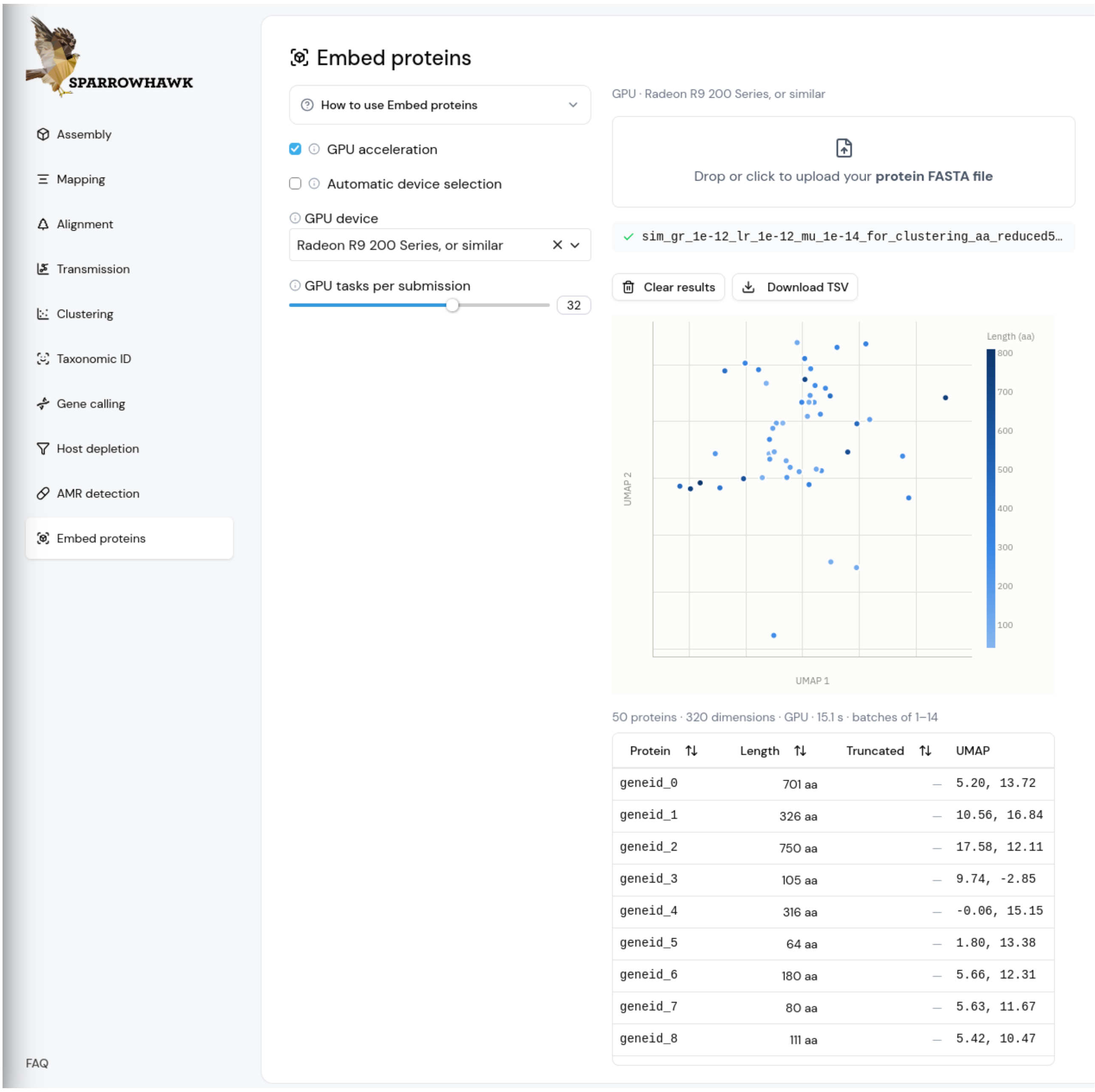


**Supplementary fig. 30.** Screenshot of `sparrowhawk` on the "Embed proteins" tab.

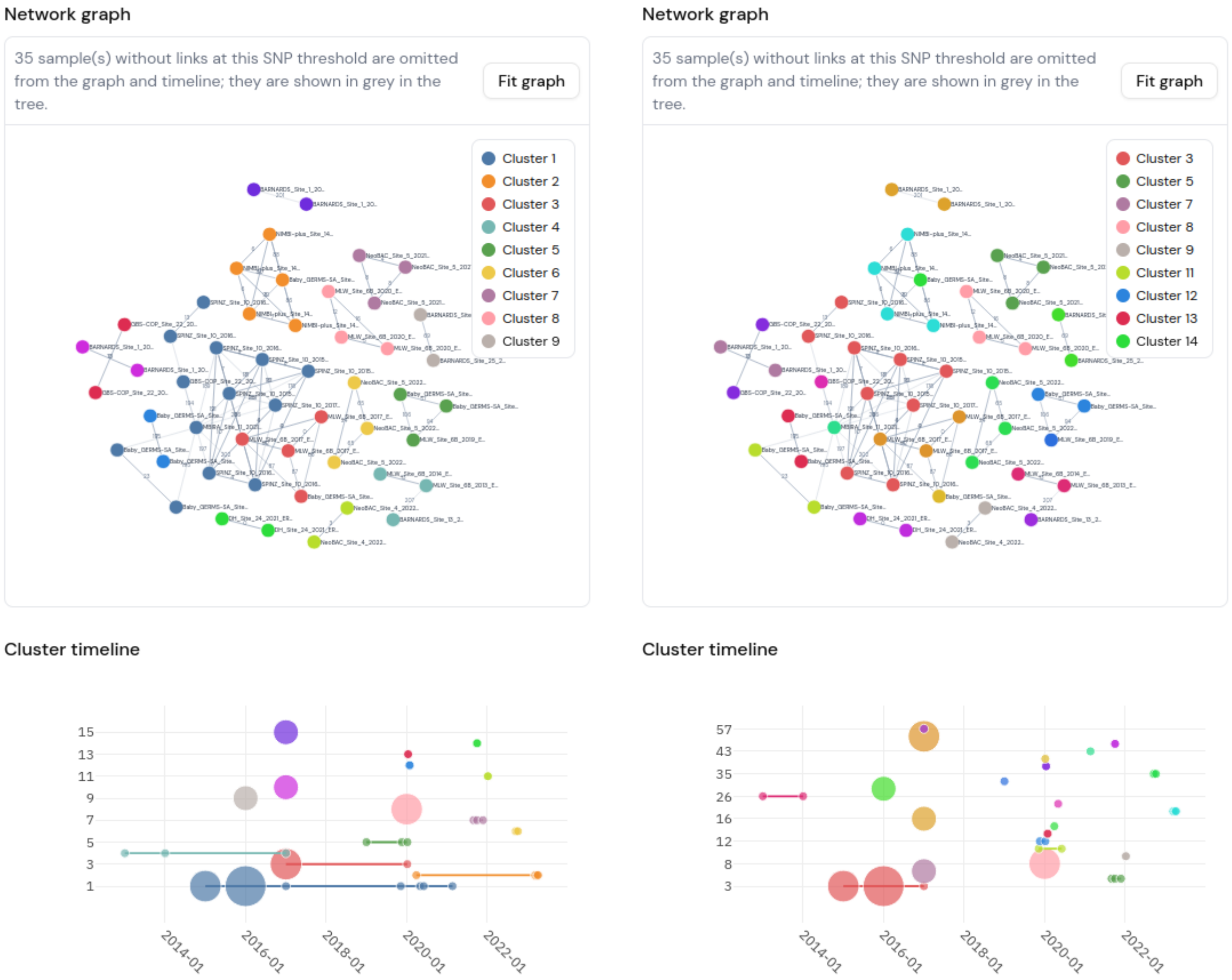


**(a)** **(b)**

**Supplementary fig. 31.** Example of how the location match affects results using the same test data on transmission results. When enabled, it separates in different clusters per location those previously identified clusters (only by single nucleotide polymorphisms, SNPs, differences). This is seen here, where the results with the "Enable location matching" checkbox activated show that the inferred reach of some SNP clusters has been broken because they are now considered two (or more) separate ones. Additionally, the network graph shows different cluster assignments to some nodes that are connected to each other. **a,** Transmission results without the "Enable location matching" box selected; **b,** with it selected.

### Supplementary tables

**Supplementary table 1.** Real-read sets of the `sparrowhawk-asm` benchmark, with thirty sequencing run accessions per species. Those marked with * were also subsampled to 5, 10, 15, and 20× coverage for the low-coverage benchmark. The list is also in `supp_asm_realreads.tsv`.

| Species | Run accessions | | | | |
|---|---|---|---|---|---|
| *Escherichia coli* | DRR567756* | DRR567759* | ERR10226929* | ERR10391689 | ERR10766880 |
| | ERR11798009 | ERR13600938 | ERR13600943 | ERR13601201 | ERR14129435 |
| | SRR13834345 | SRR29428861 | SRR29428864 | SRR29428876 | SRR30710565 |
| | SRR31106805 | SRR31348763 | SRR31612436 | SRR31612438 | SRR31612886 |
| | SRR31612892 | SRR32234844 | SRR32234858 | SRR32268898 | SRR32268902 |
| | SRR32272992 | SRR32272999 | SRR32301867 | SRR36271673 | SRR37397280 |
| *Mycobacterium tuberculosis* | ERR10680100* | ERR11082872* | ERR12311948* | SRR29318486 | SRR32565152 |
| | SRR32565155 | SRR32565156 | SRR32565159 | SRR32565164 | SRR32565169 |
| | SRR32565172 | SRR32565174 | SRR32565180 | SRR32565182 | SRR32565186 |
| | SRR32565189 | SRR32565194 | SRR32565201 | SRR32565207 | SRR32565211 |
| | SRR32565217 | SRR32565231 | SRR32565237 | SRR32565238 | SRR32565240 |
| | SRR32565242 | SRR32565243 | SRR32565244 | SRR32565246 | SRR32565249 |
| *Neisseria meningitidis* | ERR10211054* | ERR10211226* | ERR10211266* | ERR10211273 | ERR10211281 |
| | ERR10211312 | ERR10211330 | ERR10211361 | ERR10211376 | ERR10211417 |
| | ERR10211444 | ERR12055097 | ERR12055119 | ERR12055164 | ERR13147396 |
| | ERR5106357 | ERR6226274 | SRR10610817 | SRR10610857 | SRR10611048 |
| | SRR11455038 | SRR19666476 | SRR24718014 | SRR29505865 | SRR30181495 |
| | SRR30181504 | SRR32500923 | SRR32500936 | SRR32500942 | SRR35855734 |
| *Staphylococcus aureus* | DRR628060* | SRR31658480* | SRR31658556* | SRR31855147 | SRR31855274 |
| | SRR31855470 | SRR32589280 | SRR32589439 | SRR32589495 | SRR32589716 |
| | SRR32589723 | SRR32589764 | SRR32589788 | SRR32589840 | SRR32589863 |
| | SRR32589871 | SRR32589892 | SRR32589912 | SRR32595726 | SRR32595778 |
| | SRR32595826 | SRR32595922 | SRR32595937 | SRR32595982 | SRR32595997 |
| | SRR32596054 | SRR32596119 | SRR32596203 | SRR32596219 | SRR32596441 |
| *Streptococcus pneumoniae* | ERR14366219* | SRR26931854* | SRR28343502* | SRR28348682 | SRR28348687 |
| | SRR30419386 | SRR31228862 | SRR31228867 | SRR31228871 | SRR31228872 |
| | SRR31228874 | SRR31228887 | SRR31228897 | SRR31228901 | SRR31228915 |
| | SRR31228922 | SRR31228926 | SRR31228939 | SRR31228944 | SRR31228948 |
| | SRR31364749 | SRR31364758 | SRR31364760 | SRR31364773 | SRR31364774 |
| | SRR31364781 | SRR31364785 | SRR31364791 | SRR31364799 | SRR31364804 |
| *Streptomyces coelicolor* | SRR23433654* | SRR23433655* | SRR23433659* | SRR23433662 | SRR23433667 |
| | SRR23433668 | SRR23433669 | SRR23433670 | SRR24425354 | SRR24425358 |
| | SRR24425359 | SRR24425379 | SRR24425380 | SRR24425382 | SRR24425383 |
| | SRR27605478 | SRR31350266 | SRR34948729 | SRR35948870 | SRR35948872 |
| | SRR35948874 | SRR35948875 | SRR35948876 | SRR35948878 | SRR35948880 |
| | SRR35948881 | SRR35948882 | SRR35948883 | SRR35948884 | SRR35948885 |

**Supplementary table 2.** Reference genomes of the simulated read sets of the `sparrowhawk-asm` benchmark. From each reference, thirty paired-end read sets were simulated with `ART` and other thirty with `Mason`, one per random seed (the same seeds for every species and simulator), with 150 bp reads, a mean fragment length of 350 bp (standard deviation 30 bp), and 30× coverage. The list is also in `supp_asm_simulation_references.tsv`.

| Species | Strain | Assembly | Length (bp) | GC (%) |
|---|---|---|---|---|
| *Escherichia coli* | K-12 MG1655 | GCF_000005845.2 | 4,641,652 | 50.79 |
| *Mycobacterium tuberculosis* | H37Rv | GCF_000195955.2 | 4,411,532 | 65.61 |
| *Neisseria meningitidis* | MC58 | GCF_000008805.1 | 2,272,360 | 51.53 |
| *Staphylococcus aureus* | Mu50 | GCF_000009665.1 | 2,903,636 | 32.84 |
| *Streptococcus pneumoniae* | R6 | GCF_000007045.1 | 2,038,615 | 39.72 |
| *Streptomyces coelicolor* | A3(2) | GCF_000203835.1 | 9,054,847 | 72.00 |

**Supplementary table 3.** *Klebsiella pneumoniae* isolates of the transmission analysis, per study. They are the isolates of Table S3 of [1] except BG333 (Baby GERMS-SA), whose assembly could not be downloaded from ENA. The read sets were assembled with `sparrowhawk-asm` with its default settings. For Baby GERMS-SA, the published assemblies were used. The isolates, with their BioSample and run or assembly accessions, are in `supp_transmission_isolates.tsv`.

| Study | Countries | Sites | Years | Read sets | Assemblies | BioProject |
|---|---|---|---|---|---|---|
| BARNARDS | 6 | 7 | 2016–2017 | 218 | 0 | PRJEB33565 |
| Baby GERMS-SA | 1 | 6 | 2019–2020 | 0 | 139 | PRJNA796486, PRJNA1282934 |
| DH | 1 | 1 | 2019–2022 | 34 | 0 | PRJEB70311 |
| GBS-COP | 1 | 1 | 2019–2020 | 49 | 0 | PRJNA1175467 |
| KWTRP | 1 | 1 | 2013–2023 | 167 | 0 | PRJNA1265413 |
| MBIRA | 6 | 6 | 2020–2021 | 173 | 0 | PRJNA1274034 |
| MLW | 1 | 1 | 2013–2020 | 362 | 0 | PRJEB42462 |
| NIMBI-plus | 1 | 1 | 2022–2023 | 29 | 0 | PRJNA1271068 |
| NeoBAC | 1 | 2 | 2021–2022 | 66 | 0 | PRJNA1265413 |
| SPINZ | 1 | 1 | 2015–2017 | 285 | 0 | PRJEB46513 |
| Total | 13 | 27 | 2013–2023 | 1,383 | 139 | |

**Supplementary table 4.** Species of the 10,000 bacterial assemblies of the `sparrowhawk-amr` benchmark. They were selected from the 27,931 assemblies with AMR records in the EBI AMR Portal, stratified so that every ESKAPEE species and antibiotic class had at least 300 assemblies where available. AMRFinderPlus 4.2.7 (database 2026-05-15.1) calls on each assembly were the reference. The assembly and BioSample accessions are in `supp_amr_assemblies.tsv`.

| Species | Assemblies |
|---|---|
| *Salmonella enterica* | 2,971 |
| *Escherichia coli* | 1,655 |
| *Streptococcus pneumoniae* | 982 |
| *Shigella sonnei* | 858 |
| *Staphylococcus aureus* | 558 |
| *Klebsiella pneumoniae* | 491 |
| *Neisseria gonorrhoeae* | 449 |
| *Acinetobacter baumannii* | 328 |
| *Enterococcus faecium* | 327 |
| *Shigella flexneri* | 302 |
| *Campylobacter coli* | 300 |
| *Campylobacter jejuni* | 300 |
| *Pseudomonas aeruginosa* | 292 |
| *Enterobacter cloacae* | 48 |
| *Clostridioides difficile* | 35 |
| *Enterobacter hormaechei* | 19 |
| *Serratia marcescens* | 15 |
| *Proteus mirabilis* | 14 |
| *Shigella boydii* | 9 |
| *Morganella morganii* | 6 |
| *Enterobacter kobei* | 5 |
| *Enterobacter roggenkampii* | 5 |
| *Providencia rettgeri* | 5 |
| *Salmonella infantis* | 5 |
| *Shigella dysenteriae* | 5 |
| *Enterobacter asburiae* | 4 |
| *Enterobacter* sp. | 3 |
| *Shigella* sp. | 2 |
| *Enterobacter aerogenes* | 1 |
| *Enterobacter bugandensis* | 1 |
| *Enterobacter chengduensis* | 1 |
| *Proteus vulgaris* | 1 |
| *Providencia* sp. | 1 |
| *Providencia stuartii* | 1 |
| *Serratia liquefaciens* | 1 |
| Total | 10,000 |

### Supplementary notes

#### Supplementary note 1: $k$-mer hash collisions in `sparrowhawk-asm`

The `ntHash2` rolling hash function [2] is used in `sparrowhawk-asm` to hash $k$-mers and to build the de Bruijn graph that is key to the assembly algorithm of the method. During `sparrowhawk-asm`'s development, we checked for hash collisions, as they can affect the graph structure and thus the output contigs by creating spurious links between $k$-mers.

We discovered that, when processing big bacterial genomes such as *Streptomyces avermitilis*, we did have collisions in the outputs for $k > 64$ onwards, $\approx 10^4 - 10^5$ collisions. We reviewed our implementation and found that it was correct.

After some analysis, we thought this might be related to what prompted the development of `ntHash2` from its previous version, which was an inherent periodicity in the hashes produced. This issue was originated on rotations that were made to the whole hash's 64 bits. `ntHash2` fixed that periodicity by splitting the hashes in various independent parts of different lengths to which rotations were applied independently. This allowed a higher period (the least common multiple of the lengths), that in the default `ntHash2` is 1023, with the hash being splitted in two parts of 31 and 33 bits.

We explored increasing the number of splits to see if this reduced the number of collisions when $k$-mer diversity is high (as with these large genomes), by comparing what was described in the supplementary material of [2]. We found that collisions are significantly reduced with more splits. We finally decided to keep six splits of 5, 7, 9, 11, 13, and 19 bits, offering a much larger period of 855855, as this reduced collisions to a minimum $(0 - 4)$ even with the largest genomes we tried.
